# *Sphagnum* peat moss thermotolerance is modulated by the microbiome

**DOI:** 10.1101/2020.08.21.259184

**Authors:** Alyssa A. Carrell, Travis J. Lawrence, Kristine Grace M. Cabugao, Dana L. Carper, Dale A. Pelletier, Sara Jawdy, Jane Grimwood, Jeremy Schmutz, Paul J. Hanson, A. Jonathan Shaw, David J. Weston

## Abstract

*Sphagnum* peat mosses is a major genus that is common to peatland ecosystems, where the species contribute to key biogeochemical processes including the uptake and long-term storage of atmospheric carbon. Warming threatens *Sphagnum* mosses and the peatland ecosystems in which they reside, potentially affecting the fate of vast global carbon stores. The competitive success of *Sphagnum* species is attributed in part to their symbiotic interactions with microbial associates. These microbes have the potential to rapidly respond to environmental change, thereby helping their host plants survive under changing environmental conditions. To investigate the importance of microbiome thermal origin on host plant thermotolerance, we mechanically separated the microbiome from *Sphagnum* plants residing in a whole-ecosystem warming study, transferred the component microbes to germ-free plants, and exposed the new hosts to temperature stress. Although warming decreased plant photosynthesis and growth in germ-free plants, the addition of a microbiome from a thermal origin that matched the experimental temperature completely restored plants to their pre-warming growth rates. Metagenome and metatranscriptome analyses revealed that warming altered microbial community structure, including the composition of key cyanobacteria symbionts, in a manner that induced the plant heat shock response, especially the Hsp70 family and jasmonic acid production. The plant heat shock response could be induced even without warming, suggesting that the warming-origin microbiome provided the host plant with thermal preconditioning. Together, our findings show that the microbiome can transmit thermotolerant phenotypes to host plants, providing a valuable strategy for rapidly responding to environmental change.

## Introduction

*Sphagnum* peat mosses are fundamental ecosystem engineers (1, 2), contributing to the construction of bog and peatland systems that occupy just 3% of the global land surface yet store approximately 30% of all soil carbon (3, 4). In boreal regions, *Sphagnum* production can increase with modest warming (5, 6), but these positive effects are not entirely generalizable (7) and are expected to be offset by water stress from surface drying (8) and more extreme warming events (9–11). The competitive success and productivity of this keystone genus is largely dependent on symbiotic interactions with microbial associates (12–14), through which *ca.* 35% of atmospheric nitrogen fixed by diazotrophic bacteria in the microbiome is transferred to the *Sphagnum* host (15). Currently, however, we lack a basic understanding of how warming influences *Sphagnum*-microbiome interactions and how these interactions influence host acclimation and adaptation to elevated temperature.

*Sphagnum* symbiosis is characterized by an intimate association with dinitrogen (N_2_)-fixing cyanobacteria on the host cell surface and within water-filled hyaline cells (16–19). Hyaline cells provide a key function for nonvascular mosses, which are incapable of active water transport, and also provide a buffered environment for the microbiome that is less harsh than the external pore water, which is characterized by fluctuating temperature spikes and low pH (2). Phylogenetic evidence suggests that bacterial methanotrophs are also important N_2_-fixing members of the *Sphagnum* microbiome in boreal peat bogs (20–23). These methanotrophs not only fix N_2_ but also supply 5–20% of the CO_2_ necessary for host photosynthesis as a by-product of methane oxidation (24). In addition to the prominent N_2_-fixing bacteria, *Sphagnum spp.* host a diverse array of heterotrophic bacteria, archaea (12), fungi (12), protists (25, 26), and viral symbionts (27) within a complex food web structure. Results from a whole ecosystem peatland warming experiment indicates that elevated temperatures are associated with changes in the *Sphagnum* microbial community, reduced N_2_ fixation (28) and reduced *Sphagnum* biomass production (11). It remains unknown whether the warming-altered microbiome influences host acclimation, growth, and production, and if so, in what manner.

Disentangling the effects of *Sphagnum* symbiotic interactions in the context of climate change is thwarted by our inability to predict whether and how mutually beneficial interactions will persist under variable environments. In N_2_-fixing legumes (29) and coral systems (30–33), for example; altered environmental conditions can increase the cost of the interaction relative to the benefits (i.e., the cost:benefit ratio), resulting in breakdown of mutualism and even to antagonistic interactions. One strategy for maintaining a favorable cost:benefit ratio is partner switching, i.e., the substitution of one symbiont for another. In corals, for example, the negative effect of elevated temperatures on host performance can be tempered by replacing symbiont partners with more thermotolerant species (32, 33). By contrast, the habitat-adapted symbiosis paradigm does not emphasize partner choice, but instead proposes that endophytes adapt to stress in a habitat-specific manner and can confer the same functional stress tolerance to their plant hosts (34). Because it is not known whether endophytes are locally adapted or differentiated by environmental sorting, the term “adaptation” is applied loosely (35). Nonetheless, habitat-associated benefits from endophytes originating from extreme temperatures and salinities can benefit host plants subjected to the same environmental extremes (36). By contrast, the habitat origin of fungal endophytes along a rainfall gradient has little effect on the drought responses of *Panicum virgatum* (37). A more explicit test of habitat-associated effects relative to evolutionary history and physiological traits was carried out by Giauque and colleagues (35). They found little support for the idea that fungal endophyte phylogenetic relatedness predicts host benefits, but did find some evidence that microbes that had experienced similarly stressful environments could benefit their hosts. However, the host benefit was not as strong as in previous studies (34), in which fungal endophytes were isolated from more extreme environments. Further complicating the habitat-adaptation paradigm is our lack of understanding of the underlying mechanism or the roles of non-fungal members in conferring host benefits.

Given the importance of bacteria for *Sphagnum* performance and ecosystem biogeochemistry (12, 14, 23, 24), we sought to determine the influence of habitat origin on host acclimation to thermal stress. To investigate this experimentally, we mechanically separated the microbiome from field-grown *Sphagnum* plants collected under two extreme thermal conditions, transferred the constituent microbes to germ-free plants, and then exposed the new host plants to short-term heat stress. To assess host and bacterial dynamics, we performed growth analysis, chlorophyll-*a* fluorescence imaging, metagenomics, metatranscriptomics, and 16S rDNA profiling. The transfer of environmentally conditioned microbiomes to germ-free plants, which is analogous to microbiome transplant studies in medical research, will allow us to (i) determine whether a warming-conditioned microbiome can transmit thermotolerance to the plant host, (ii) characterize the community structure and relative species abundance of beneficial microbiomes, and (iii) allow in-depth exploration of key genes meditating microbially induced host thermotolerance.

## Results

**Fig. 1.**
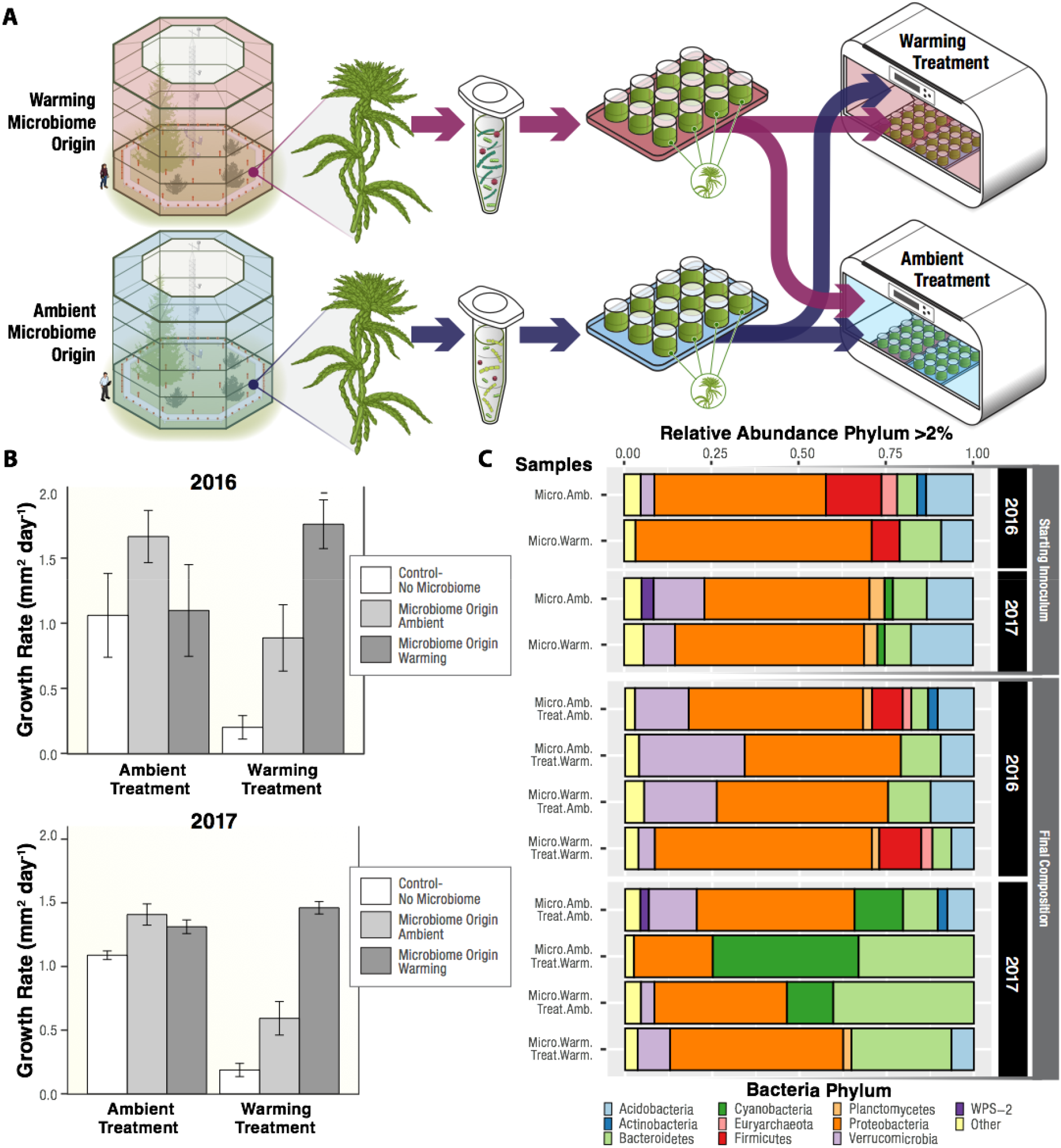
Experimental approach and design: Field-collected donor moss microbiomes collected from ambient or warming conditions were transferred to germ-free recipient moss, and the resulting communities were then placed in an ambient or warm growth chamber (A). Average moss growth rate under ambient or warming treatments, as a function of the thermal origin of the microbiome. Error bars represent standard error of the mean of n = 6 for 2016, n = 12 for 2017 (B). Relative abundance of microbiome phyla, determined by 16S rDNA amplicon sequencing of the starting field-collected inoculum from ambient or warming experimental plots, and the final compositions of experimental samples (C).

### Plant host performance in response to experimental temperature is dependent on the thermal origin of the microbiome

In a previous study (28) conducted at a whole-ecosystem peatland manipulation experiment (SPRUCE (38)), we discovered that *Sphagnum*-associated microbiome composition and N_2_ fixation were influenced by above- and below-ground warming treatments. To determine the influence of microbiomes from contrasting thermal habitats on plant host thermal acclimation and performance, we conducted microbiome transfer experiments. Starting inocula were isolated from field plants in which we mechanically separated microbes from their *Sphagnum* hosts collected at the SPRUCE experimental field plots exposed to ambient + 0°C (referred to as the ambient microbiome origin) or ambient + 9°C (warming microbiome origin) temperatures Fig. 1A). Independent replicates were collected in two consecutive years (2016 and 2017). Treatments included microbiomes collected from ambient and warming origin conditions, along with a microbiome-free mock control. After transfer of field inocula to laboratory-grown *S. angustifolium,* the constructed communities were exposed to warming or ambient temperature conditions in a full factorial design (Fig. 1A). The repeated 2017 experiment used the same experimental design, except that the number of replicate plants was increased from *n* = 6 to *n* = 12. The microbiome thermal origins were similar to conditions in the experimental chambers (*SI Appendix, Table S1*).

In both years, a donor microbiome that matched the experimental thermal conditions conferred the greatest increase in host growth (Fig. 1B). Host benefits from the microbiome were especially apparent under experimental warming conditions: in 2016 and 2017, moss receiving a warming-origin microbiome exhibited an increase in growth of 87% and 89%, respectively, relative to plants receiving the mock control (Fig. 1B; *SI Appendix, Tables 2–4*). The addition of a discordant (temperature-mismatched) microbiome increased moss host growth relative to the control by 77% and 68% in the 2016 and 2017 experiments, respectively, although post-hoc tests were not significant within year at a *P* < 0.05 alpha level. Generally, benefits to plant host were most pronounced under experimental warming conditions, under which plants without microbes were severely affected (Fig. 1B). Host benefits from temperature-mismatched microbiomes were least prominent under ambient experimental treatment, in which growth increases ranged from 3 to 17% following inclusion of a warming-origin microbiome (*F*_temperature:microbiome_ = 32.01; *P* < 0.05 for 2017; *SI Appendix, Table 3*). Throughout the experiment, moss photosynthetic activity and response to temperature and microbiome origin were evaluated by monitoring chlorophyll-*a* fluorescence (*F*_v_/*F*_m_). The results mirrored the growth analysis: *F*_v_/*F*_m_ values were higher when microbiome thermal origin matched experimental temperature (*SI Appendix, Table S5-6 & Fig. S1–2*). Thus, the warming microbiome consistently increased moss performance to warming, as reflected by growth rate and photosynthetic activity, demonstrating that microbiome thermal origin can play a substantial role in host moss acclimation to elevated temperatures.

### Habitat origin and thermal treatment conditions structure the starting microbiome and resultant microbial community

**Fig. 2.**
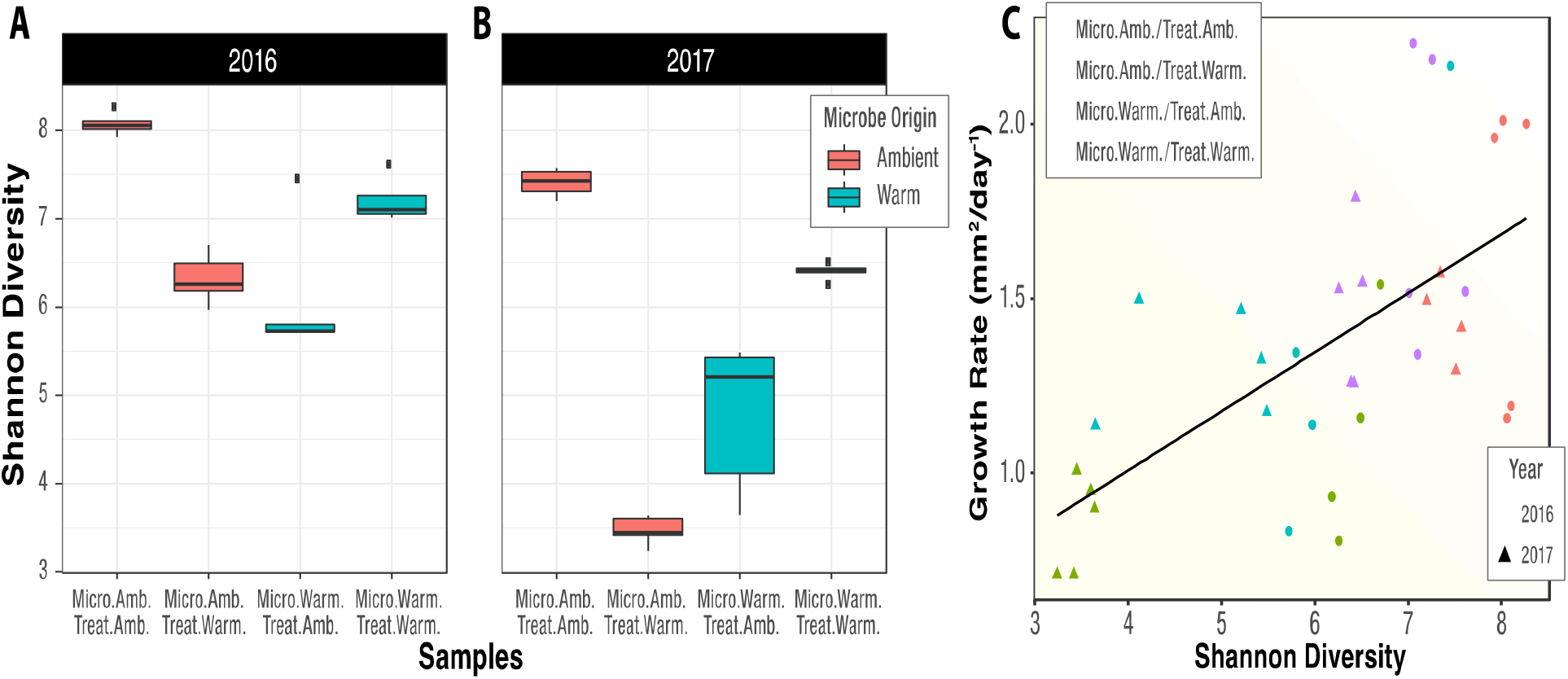
Shannon diversity index of the microbiome at the conclusion of the experiments in 2016 (A) and 2017 (B), based on 16S rDNA amplicon data. Microbiomes were less diverse when the thermal origin and experimental treatment were mismatched (i.e., Ambient Origin in Warming Treatment or Warming Origin in Ambient Treatment). Plant growth rate was linearly correlated with microbiome Shannon diversity at the conclusion of the experiment (C).

In 2017, the initial community structure of the *S. angustifolium* field-collected inoculum differed between thermal origins (Adonis, R^2^ = 0.92133, *P* = 0.009) and 2016 (Adonis, R^2^ = 0.53, *P* = 0.1) (*SI Appendix, Fig S3*). At the class level, the ambient-origin microbiome consisted largely of Alphaproteobacteria (32%) and Clostridia (16%) in 2016, whereas Alphaproteobacteria (30%) and Acidobacteria (11%) were most abundant in 2017 (*SI Appendix, Table S7*). Within the warming-origin microbiome, Gammaproteobacteria were highly abundant in 2016 (43%), but only constituted 15% of community abundance in 2017, with the difference compensated by an additional increase in Alphaproteobacteria abundance (30%). Despite between-year differences in community composition at the class level within thermal regimes, the growth benefits provided to the plant were strikingly consistent (Fig. 1B).

*Sphagnum*-associated microbial communities responded to four weeks of thermal treatment conditions regardless of year, thermal origin, or growth temperature. Non-metric multidimensional scaling (NMDS) ordinations of the microbiome Bray-Curtis distance matrix revealed that the community composition of the warming-origin microbiomes responded similarly across thermal treatments, whereas ambient-origin microbiome structure varied to a greater extent both across and within thermal treatments (*SI Appendix, Fig S4*). To determine whether changing community composition influenced microbial diversity, we estimated the Shannon diversity index for each treatment condition at the conclusion of the study in both years (Fig. 2A–B). Microbial diversity was highest when microbiome thermal origin matched chamber treatment temperatures; conversely, discordant combinations resulted in substantially lower microbial diversity (Fig. 2A–B). The ambient-origin microbiome had the highest diversity under matched (i.e., ambient) treatment conditions in both years (ANOVA, *P* < 0.01). Similarly, the warming-origin microbiome had the highest diversity under the warming treatment in both years (ANOVA, *P* < 0.01). Detailed class-level community composition assignments are provided in *SI Appendix, Table S8.* Given that greater phylogenetic diversity is likely to be accompanied with greater metabolic and functional diversity, we hypothesized that microbial diversity would be associated with enhancements in plant acclimation to stressful warming conditions, reflected by improved growth. Providing correlative support for this hypothesis, bacterial and archaea diversity (as inferred from 16S rDNA) at the end of the experiment was correlated with *Sphagnum* growth (Pearson correlation, r = 0.744, p=0.003; Fig.2B). By contrast, ITS-derived fungal diversity estimates did not correlate with moss growth (Pearson corr., r = – 0.204, *P* = 0.403; *SI Appendix, Fig S5*), and indeed the fungal communities did not vary greatly across treatments (*SI Appendix, Table S9-10*), therefore we largely focused on the bacterial component of the microbiome in proceeding sections.

### Metagenome and metatranscriptome analyses reveal changes in symbiotic *Cyanobacteria* abundance and composition, and host plant transcriptional reprogramming in response to temperature treatment

**Fig. 3.**
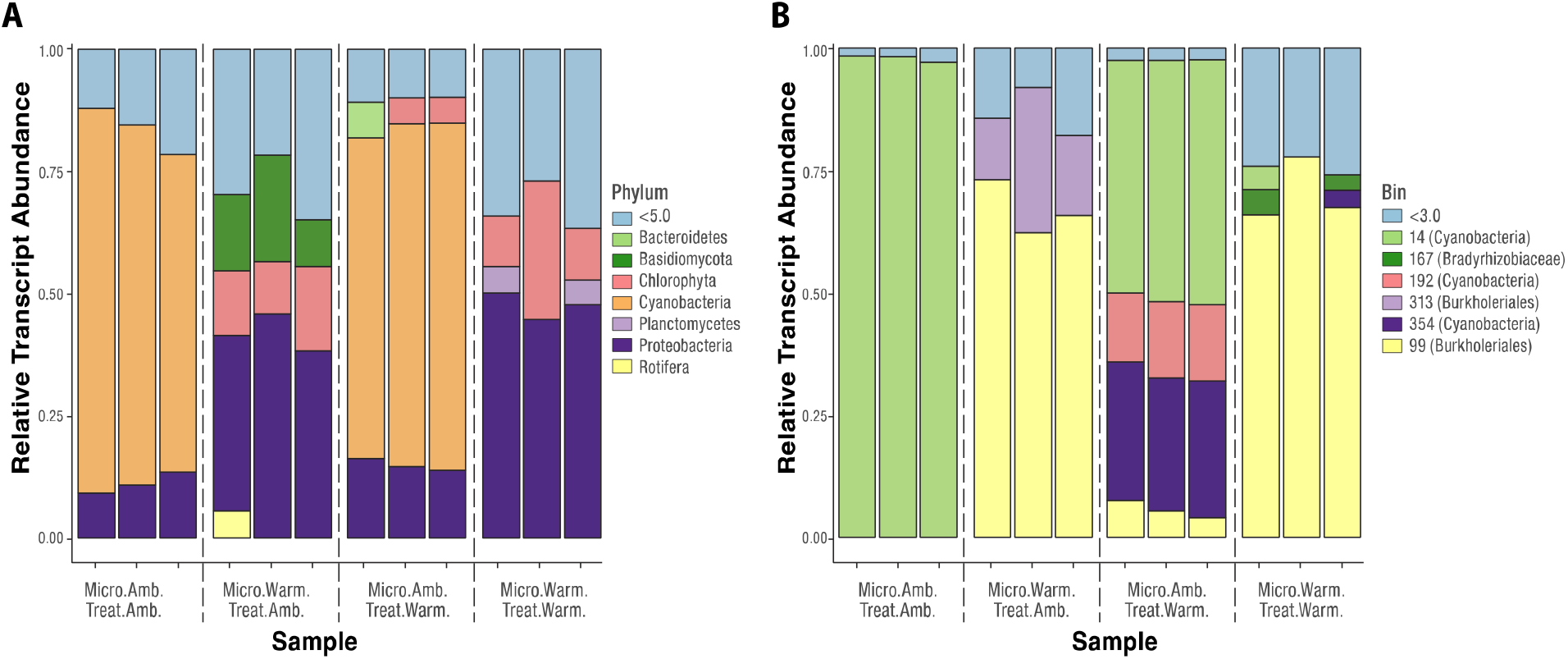
Relative abundance of microbial transcripts mapping to metagenome contigs for (A) major phyla and (B) metagenome-assembled genomes (MAGs). Each bar represents a metatranscriptome sample for the ambient-origin (Micro.Amb.) or warming-origin (Micro.Warm) microbiome under either the ambient (Treat.Amb.) or warming (Treat.Warm) treatment. Colors indicate (A) phyla or (B) MAGs; light blue represents (A) phyla with <5% or (B) MAGs with <3% of mapped transcripts.

Host thermal acclimation and productivity varied with microbiome origin. To further explore how community dynamics influence host thermal acclimation, we used metagenomics and metatranscriptomics to identify both plant and microbial gene sets responsive to thermal and microbiome conditions (Fig. 3). For metagenome assemblies, DNA sequencing reads mapping to the *S. angustifolium* genome (https://phytozome-next.jgi.doe.gov/info/Sfallax_v1_1) were removed, and the remaining reads were co-assembled into 4,762,069 contigs with an N50 of 1261 bp. Binning of metagenome contigs yielded 45 metagenome-assembled genomes (MAGs) with a quality score ≥ 70 with ≤ 5% contamination (*SI Appendix Supplemental Table S11).* The high-quality MAG standard of >90% complete and <5% contamination (39) was met for 28 of our genomes, whereas 13 and 9 MAGs are >95% and >97% complete, respectively. Taxonomic assignments and blast hits from annotated proteins were resolved to the lowest taxonomic level using CheckM (40) and DIAMOND (41) (*SI Appendix, Table. S11*). For metatranscriptomes, we generated 429.6 GB of RNA-seq data across three replicates for each treatment. On average, 40.93 million (M) ± 7.39 M reads passed quality filtering per sample across all thermal treatments and microbiome conditions (*SI Appendix,* Table. S12). In samples derived from plants receiving a microbiome transfer, approximately 65% of reads aligned to the *Sphagnum* genome, except in the discordant case when plants received ambient-origin microbiomes followed by warming treatment. Under that condition, the plants were severely stressed, and only 12.4% of the reads aligned to the *Sphagnum* genome (*SI Appendix,* Fig. S6).

To expand on the amplicon-based community composition results (Fig. 1C, *SI Appendix,* Table S7) and determine which microbial members are transcriptionally active, we categorized transcriptional profiles based on taxonomic composition. Under matched ambient origin and experimental temperature, microbial transcripts were mostly from Cyanobacteria symbionts (72.5 ± 6.9%), followed by Proteobacteria (11.2±2.2%) (Fig. 3A). Under matched warming origin and temperature treatment, Cyanobacteria transcript reads were largely absent, and the metatranscriptome was mainly derived from Proteobacteria (47.5±2.7%), Chlorophyta (16.39±10.4%), and Planctomycetes (4.9±0.57%). Results from mismatched origin and experimental conditions more closely reflected their microbiome origin communities (*SI Appendix, Table S7*). This finding was also reflected in a multidimensional scaling analysis using level 3 SEED functional annotation, in which cluster variation was explained more on microbial origin rather than experimental temperature (*SI Appendix, Fig. S7*).

### Experimental warming increases transcript abundance from alternative cyanobacteria members, signaling possible symbiont exchange

**Fig 4.**
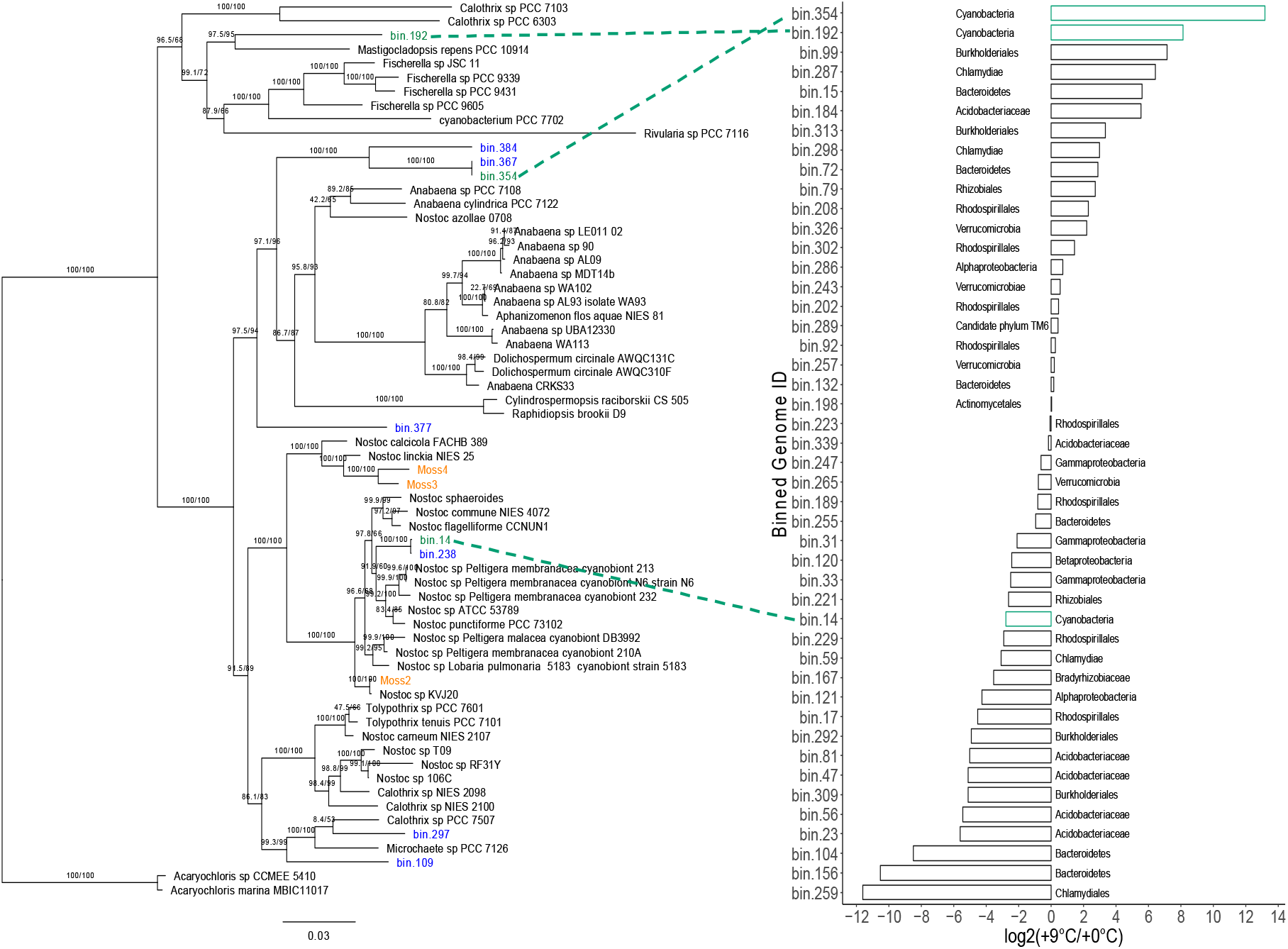
Phylogeny of select Cyanobacteria and log_2_(fold change) of metagenomic bins. (A) Maximum-likelihood phylogram where the numbers at nodes indicate UFBoot2 and SH-like approximate likelihood ratio support. Branch lengths indicate estimated substitutions per site. Metagenomic bin taxa labels are colored by source within SPRUCE enclosures (green), outside SPRUCE enclosures (blue), or from (42) (orange). (B) Bar chart representing log_2_(fold change) of metagenomic bins between ambient-origin and warming origin metagenomes. Green bars indicate cyanobacterial MAGs recovered in this work.

Differences in the composition and abundance of *Sphagnum*-associated cyanobacteria in response to warming are important because these organisms are key symbionts with *Sphagnum* mosses, and the exchange of symbionts with more thermotolerant forms has been implicated in host thermotolerance in coral systems (e.g., 34). To explore this further, we taxonomically refined three of our high-quality cyanobacteria MAGs with a phylogenetic tree reconstruction using an additional 109 cyanobacterial genomes (Fig. 5). We found that all three cyanobacterial MAGs belong to the heterocystous B1 clade of Cyanobacteria, which also contains known plant associates (43). To determine which of the cyanobacteria are most responsive to thermal conditions, we aligned the microbial RNA-seq reads from the end of the experiment onto the MAGs. Of all *non-Sphagnum* RNA-seq reads, 31.6±8.7% mapped onto cyanobacteria MAGs for matched ambient-origin and ambient temperature conditions. This percentage decreased to 26.1±1.1% when plants receiving ambient-origin microbiomes were subjected to discordant warming treatment. Further, microbiomes originating from warming field conditions contained negligible levels of cyanobacterial RNA-seq reads (0.1–0.08%). Cyanobacterial reads predominantly aligned to MAG bin 14 (98±0.7%), but to a considerably lesser extent (49±0.6%) when placed under warming experimental conditions. The decrease in bin 14 RNA-seq reads was accompanied by an increase in reads from *Cyanobacteria* bins 354 (28±0.6%) and bin 192 (15±0.9%) (Fig. 3B). Due to sampling constraints, we did not normalize the results of RNA-seq analysis to community abundance changes. Nonetheless, in light of abundance estimates based on metagenomic data (Fig. 3), field-collected 16S rDNA amplicon results from this study (Fig. 1), and results of a prior study (26), we are confident that warming decreases the abundance of *Sphagnum*-associated cyanobacteria and changes cyanobacterial membership of the community.

### Host plant transcriptional reprogramming and acclimation in response to warming are consequences of microbiome thermal origin

**Fig 5.**
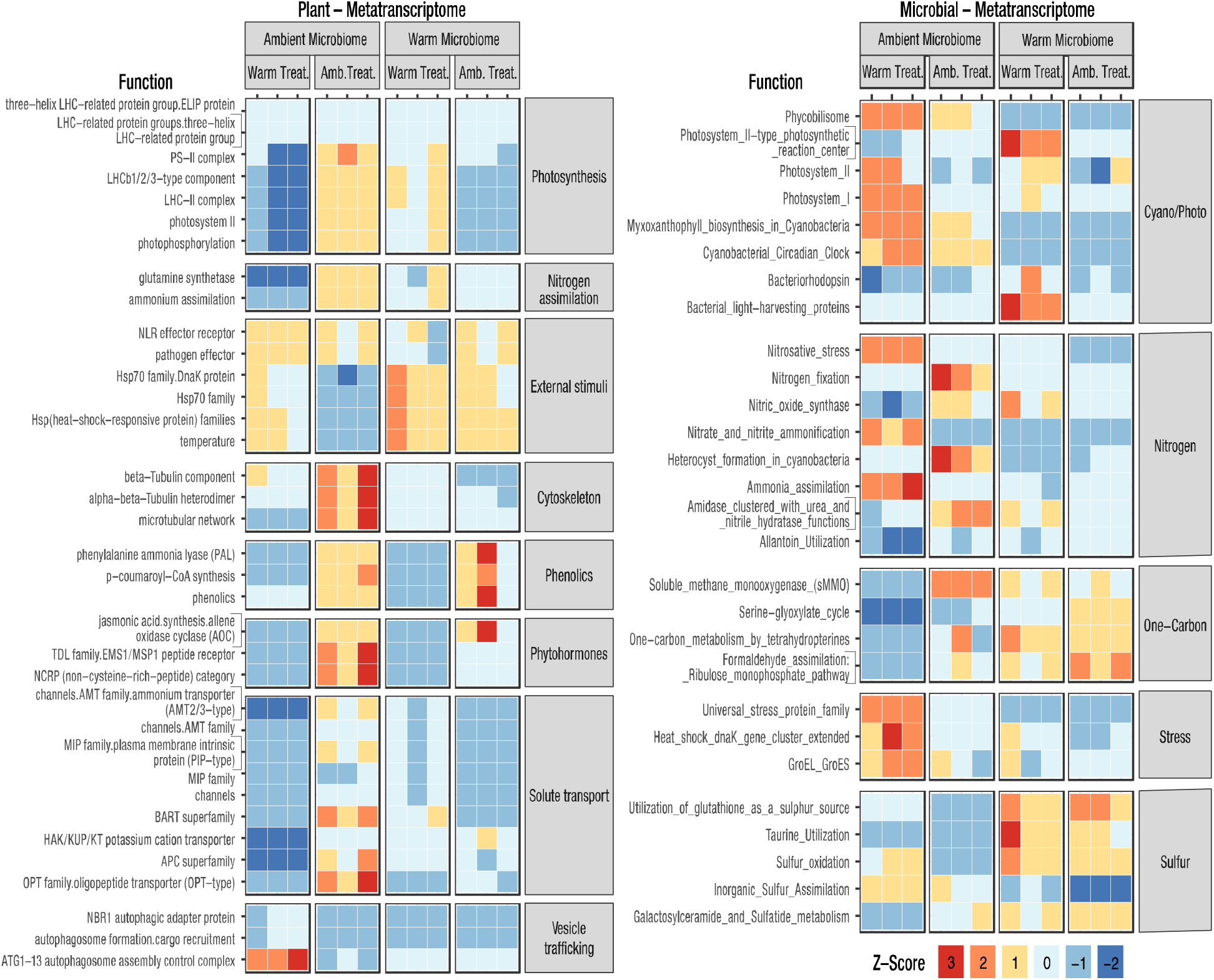
Heatmap of z-scores for (A) MapMan4 ontology categories enriched in differentially expressed plant genes and (B) differentially expressed SEED level 3 categories related to nitrogen, one-carbon, and sulfur metabolism; cyanobacterial/photosynthesis; and stress. Differential expression was defined as |log_2_(Fold Change) > 1| and corrected p < 0.05.

Given that the warming environment alters community composition in a way that benefits host plant acclimation to warming, we hypothesized that the warming environment selects for more thermotolerant symbionts that are able to maintain nitrogen exchange with the plant at elevated temperatures. If this is the case, we would expect that plant and microbial transcriptional patterns relating to N transport and metabolism would be similar between matched origin and temperature conditions (i.e., warming origin + warming treatment or ambient warming + ambient treatment). Functional ontology enrichment analysis across all conditions revealed that in plants receiving an ambient-origin microbiome and ambient treatment condition, gene expression was enriched for pathways involved in N metabolism, including ammonium transporters, ammonium assimilation, and glutamine synthetase, as well as growth-related ontologies including photosynthesis, cytoskeletal elongation, and hormonal regulation (Fig. 4A). This was also apparent on a log_2_(fold change) (LFC) basis when comparing plants with an ambient-origin microbiome between temperature conditions (*SI Appendix, Dataset S1*). In this case, ambient treatment plants corroborated enrichment analysis with induced ammonium transport (LFC 4.0, *P* = 2.0 × 10^−2^) and glutamine synthetase (LFC 2.4, *P* = 1.1 × 10^−2^). In addition, 153 of 178 genes within the photosynthesis ontology were induced, with photosystem II light-harvesting complex II most strongly affected (LFC 2.1, *P* = 2.48 × 10^−6^). In addition, we noted differences in fatty acid synthesis, especially in desaturation and elongation (LFC 2.74, *P* = 2.97 × 10^−6^), phenolic secondary metabolite production (LFC 2.86, *P* = 1.77 × 10^−5^), cell wall expansion (LFC 6.5, *P* = 2.7 × 10^−6^), phytohormone signaling with non–cysteine-rich peptides (LFC 4.63, *P* = 8.1 × 10^−4^), jasmonic acid synthesis (LFC 2.36, *P* = 4 × 10^−2^), and response to external stimuli (LFC 2.53, *P* = 3.2 × 10^−2^). As expected, heat shock proteins (HSPs) were responsive to warming, especially the HSP70 family, of which 15 members were induced (LFC 1.2, *P* = 6.5 × 10^−3^).

Plant ontology enrichment analysis did not support the hypothesis that the warming-origin microbiome would provision the plant with N at warming treatment conditions (Fig. 4A). Likewise, there was no support for this hypothesis on an LFC basis when comparing RNA-seq profiles from plants with warming-origin microbiomes across temperature treatments (*SI Appendix, Dataset S1*). Despite the apparent lack of microbially provided fixed N, the warming-origin microbiome still provided growth benefits to warming-treated plants (Fig. 1B), and this was also apparent in RNA-seq enrichment analysis of growth-related ontologies. Specifically, plants exposed to warming that received warming-origin microbes exhibited enrichment for photosynthesis – photosystem II light harvesting complex II (LFC 1.4, *P* = 1.9 × 10^−3^), cell wall expansion (LFC 2.5, *P* = 1.3 × 10^−5^), and phenolic secondary metabolite production (LFC 2.9, *P* = 2.0 × 10^−8^).

Microbial RNA-seq differential expression (DE) analysis of functional ontologies supported the notion that the warming-altered cyanobacteria community is not fixing N, and is therefore not provisioning the plant with N. DE enrichment analysis revealed that microbial N metabolism differed dramatically between both treatments and origins (Fig. 4B, *SI Appendix,* Table S13). Indeed, exposure to warming decreased N-fixation ontology gene expression by 53.4-fold (Fig. 5B, *SI Appendix,* Table S13). Moreover, there was no enrichment evidence for N-fixation among the warming-origin microbiomes, regardless of temperature treatment. Although we could not obtain direct evidence for N-fixation in this study due to sample size restrictions, these observations corroborate prior ^15^N_2_-based fixation rates reported at the same field site where our warming- and ambient-origin inocula were obtained (28).

How the warming-origin microbiome influences host plant photosynthesis and growth temperature acclimation remains to be elucidated, but we can glean clues from communities composed of discordant warming-origin microbes at ambient experimental temperatures. In that case, enrichment for the HSP70 family and phenolic compounds are induced without heat (Fig. 4A). This trend was also observed on a LFC basis where 33 out of 35 detected HSPs were induced (*SI Appendix, Dataset S1*). This suggests that the warming-origin microbiome is eliciting, or preconditioning, the host plant heat shock response without the need for elevated temperatures.

## Discussion

The establishment of constructed communities derived from microbiome transfers, coupled with comparative metatranscriptomics, revealed several novel aspects of microbial contributions to plant temperature response. First, plants receiving a microbiome from a high-temperature environment exhibited enhanced photosynthetic and acclimation responses to similarly warm environments. Second, the warming-origin microbiome was less diverse than the ambient field-collected microbiome, but contained transcripts from a more diverse set of cyanobacteria, suggesting symbiont swapping or replacement. Finally, the warming-origin microbiome transferred a thermotolerant phenotype to the plant through the induction of host genes involved in the heat shock response and hormonal regulation.

Our results demonstrate that the originating thermal habitat of the microbiome has a dramatic effect on *Sphagnum* host acclimation to elevated temperatures. These results were consistent across two years of field-collected donor inocula and two independent laboratory experiments. Although the fact that plants benefit from microbial relationships is well known, the transfer of microbially acquired habitat specific tolerance to recipient plants was reported much more recently, and to date has been limited to endophytic fungi (44). In an early example of this, Redman et al. (45) collected a native north American grass, *Dichanthelium lanuginosum,* endemic to geothermal sites with soil temperatures reaching up to 50°C. After isolation of a *Curvularia sp.* fungus and re-inoculation onto endophyte free plants, thermotolerance was conferred to the recipient plant host. This approach of isolating endophytic fungi from plants endemic to extreme habitats in an attempt to confer habitat-associated benefits has been tested a number of times with both successful (34, 36) and mixed results (35, 37). In all cases, the microbial component focused on fungi and was constrained to single member strain-based studies.

The microbiome transfer approach used in this study allowed us to test habitat-associated benefits from a broader set of organisms that is more representative of the dynamic coevolving community. However, this strategy made it difficult to relate specific taxa to recipient host benefits. For example, the warming-origin inocula differed substantially between years, even at the phylum level, yet both provided host thermal benefits. This is consistent with the idea that microbial community taxonomic composition is not necessarily a clear indication of community function. Indeed, functional similarity independent of taxonomic group has been reported in other systems, including human gut (46) and microalgae (47) microbiomes. Hence, the challenge is to look beyond taxonomic association and determine what components of the microbiome are responsible for conferring thermotolerance on the host plant.

One possible mechanism for enhanced host temperature tolerance is the replacement of primary symbionts with more thermotolerant symbionts. *Sphagnum* mosses have long been known to host N2-fixing Cyanobacteria as symbionts (16–19). More recently, they have been shown to associate with a suite of bacteria, including those that oxidize methane into CO_2_, as well as a number of viral, archaea, and protists (reviewed in (12)). The influence of warming on these symbionts, especially the Cyanobacteria, would directly affect host nutrient status and productivity. Our metagenome analysis in this study assembled three Cyanobacterial MAGs. DNA and RNA-seq reads mapping to the binned MAG genomes indicated that *Sphagnum* plants were primarily colonized by a single Cyanobacterial member from genus *Nostoc.* With increasing temperature, this *Nostoc* MAG decreased abundance, while two additional Cyanobacterial MAGs are increasing in abundance, indicating a possible exchange for more thermotolerant members of the clade. Precedent for symbiont shuffling has been provided in coral systems. Coral host algal symbiont communities that are genetically diverse and susceptible to symbiont loss due to environmental stress and ensuing coral bleaching events. However, the stress events leading to the bleaching, as well as the bleaching itself, provide an opportunity for replacement of symbionts with organisms that are more suitable to the new environmental condition, such as those with higher stress tolerance (reviewed in (48)). The coral system also demonstrates the potential role of the surrounding bacterial community in coral thermotolerance. This was elegantly demonstrated by Ziegler and colleagues (49), who showed that long-term temperature elevation modified the composition of the bacterial community, and that particular bacterial taxa could predict coral thermotolerance. However, the coral system is not amenable to germ-free host strains or microbiome transfers, making it difficult to quantify the contribution of the microbiome to host thermotolerance.

From the results of this study, it is difficult to discern whether host thermal benefits from the donor microbiome are driven by community change from primary *Nostoc* Cyanobacteria symbionts, or instead by the surrounding microbial community. Our hypothesis that the key cyanobacteria symbiont was augmented by additional thermotolerant cyanobacteria in order to maintain N_2_ fixation at elevated temperatures was not entirely supported. Although we observed an increase in cyanobacteria diversity, and thus possible evidence for exchange with thermotolerant symbionts, the metatranscriptome analysis yielded no evidence for N_2_-fixation under warming. This is consistent with a prior field study (28) at the same SPRUCE site, where 16S rDNA amplicon profiling, *nifH* qRT-PCR, and N_2_ incubation assays revealed a decrease in *nifH-*containing N-fixing bacteria and a reduction in N_2_ incorporation in response to warming.

Despite the lack of evidence for a contribution of N_2_-fixation in contributing to plant thermotolerance, the metatranscriptome analysis did reveal a role for host plant heat shock reprogramming. Plants that never received a warming treatment were enriched for Hsp70 gene family transcripts when they received a warming-origin microbiome, but not when they received an ambient-origin microbiome. The Hsp70 family is often associated with thermotolerance: multiple studies have reported that thermotolerance is decreased by Hsp70 antisense and increased by Hsp70 overexpression (50). It should be noted that our metatranscriptome analysis did not show microbial-mediated repression of the Hsp90 family or induction of the Hsp101 family, both of which have been implicated in the heat shock response. Microbially mediated repression of *Arabidopsis* Hsp90, leading to elevated thermotolerance, was previously demonstrated in the desert-dwelling fungus *Paraphaeosphaeria quadriseptata* (51).

In addition to Hsp70 reprogramming, the warming-origin microbiome elicited host plant expression of genes contributing to jasmonic acid synthesis (*via* allene oxide cyclase [AOC]). Jasmonic acid is a key phytohormone contributing to both abiotic and biotic stress responses, and has been implicated in flowering plant thermotolerance (52). AOC synthesizes 12-Oxo-phytodienoic acid (OPDA), which is a signaling compound and intermediate in the jasmonic acid biosynthesis pathway. In the liverwort *Marchantia polymorpha,* overexpression of AOC increases OPDA, suggesting that its function is similar to that of its homologs in flowering plants (53). However, AOC overexpression in *M. polymorpha* decreases growth. Likewise, the warming-origin microbiome induced expression of enzymes involved in the production of phenolic compounds, including phenylalanine ammonia lyase (PAL), which has been implicated in both temperature response and disease resistance (54). In contrast to the heat shock response, the jasmonic acid and phenolic ontology enrichments disappeared after the plants were exposed to warming. It remains to be determined whether these compounds are contributing to a beneficial thermal preconditioning or instead reflect a defensive response.

One unexpected observation was that the warming-origin microbiome elicited the induction of the heat shock response in plants that were never exposed to elevated temperatures. Thermotolerance can be acquired by prior exposure to a sublethal temperature stress (55). Similarly, plants associated with beneficial rhizosphere microbes can more rapidly mount a defense response to biotic and abiotic stressors (56). Although there is a considerable body of literature on the biotic aspects of microbial-induced plant priming, increasing evidence suggests that plants can also be primed against abiotic stressors. For example, Ali et. al (57) found that a *Pseudomonas* sp. strain isolated from pigeon pea endemic to an arid region conferred enhanced survival and growth on sorghum seedlings exposed to elevated temperatures. This early example has been corroborated with multiple plant hosts and microbial strains, yet the underlying genetic mechanisms remain to be identified (53 and citations within). In most cases studied thus far, microbial-induced resistance to abiotic stress has been studied in individual strains or small community consortia. By contrast, in this study we examined how microbial dynamics within large communities interact to influence host physiology and growth.

## Conclusions

Our findings provide a starting point for future studies that systematically decouple inherent host acclimation responses to challenging environmental conditions from those of the associated microbiome. A key benefit of the microbiome transfer and constructed community approaches described here is that it allows the coevolved host–microbiome consortia collected from extreme environmental conditions to be separated and tested across a range of thermal experimental conditions. Our observation that the microbiome can transmit thermotolerant phenotypes has a number of implications. It sets the stage for moving beyond the current notion that plants are restricted to “adapt or migrate” strategies for survival to rapidly changing environmental conditions (59). The current study provides an alternative perspective on these outcomes by showing that thermotolerant phenotypes can be rapidly transmitted to plant host. We anticipate multiple challenges as the findings of our studies are transferred beyond the laboratory into ecological systems. First, additional research is needed to determine the extent to which inter– and intra-specific genetic variation influences the plant’s ability to receive microbial benefits, and if so, identifying the causal alleles. Bringing this goal closer to reality, a genome sequencing campaign representing some 55 species within the approximately 300-member *Sphagnum* genus, as well as the development of high-density genetic maps from sequencing of a 200-member pedigree cross, are currently underway (60). Second, the identification of responsible microbial taxa is challenged by large community diversity, complex community interactions and strain isolation limitations. These experimental tests could take multiple forms, from the dilution and sequencing of donor microbiomes to strain isolation and testing in our demonstrated plate-based experimental system. Within the context of this study, such an approach could determine whether microbial benefits are mainly a function of swapping primary cyanobacteria symbionts for more thermotolerant members, or whether additional microbial members are driving the host phenotype. Finally, determining the impact of plant host-microbiome interactions within the context of competition poses a major challenge. For example, is the evolution of mutualistic interactions a key driver in *Sphagnum* versus vascular plant competition? If so, how does changing environmental conditions influence these mutualistic interactions within a competitive framework, and how does that scale to alter peatland C and nutrient cycling dynamics across regional and global scales (61, 62)?

## Materials and Methods

### Study site and field sampling

The Spruce and Peatland Responses Under Changing Environments (SPRUCE) experiment, located in the S1 bog of the Marcell Experimental Forest (47° 30.4760’ N; 93° 27.1620’ W; 418 m above mean sea level), MN, USA, employs a regression-based design at the whole-ecosystem scale to produce nominal warming of ambient +0°C, +2.25°C, +4.5°C, +6.75°C, and +9°C in a *Picea mariana–Sphagnum spp.* raised bog ecosystem with open-top chamber systems (38). Heating of the soil was initiated in June 2014 and aboveground air heating began in June 2015. A full discussion of experimental details and ecosystem description is available (38). To obtain field-conditioned microbiomes, living green stems of *Sphagnum* were collected from ambient + 0°C (ambient) and ambient + 9°C (elevated) plots in August 2016 and August 2017. The collected stem portion typically included capitula and 2-3 cm of living stem. The *Sphagnum* material was placed in sterile bags and shipped to Oak Ridge National Laboratory on blue ice.

### Isolation of microbiomes, application to gnotobiotic *Sphagnum,* and warming treatment

To isolate the microbiomes from the field samples, 100 g of tissue was diced with a sterile razor blade and pulverized in PBS with a mortar and pestle. The resulting suspension was filtered through Mira Cloth, centrifuged to pellet the microbes, and then resuspended in 500 ml BG11 -N medium (pH 5.5). A single capitulum of axenic *Sphagnum angustifolium* was added to each well of a 12-well plate and inoculated with 2 ml of ambient-origin microbiome, warming origin microbiome, or sterile media. *Sphagnum angustifolium* genotype was the same genotype that was sequenced by the DOE JGI (https://phytozome-next.jgi.doe.gov/info/Sfallax_v1_1). The plates were placed into growth chambers with a 12hr:12hr light:dark cycle, programmed to either ambient or elevated field plot temperatures. June 2016 field plot temperatures from 6-hour blocks were averaged from each day, resulting in a cycle of 4 temperatures. June 2017 temperatures did not differ from those in June 2016, so the same temperature profile was used for incubations for both years (*SI Appendix, Table 1*).

### Measurement of growth and photosynthesis

To measure growth, black and white images were collected weekly, and surface area was measured using the ImageJ software (63). Change in surface area was determined as a proxy for growth. To estimate maximal PSII quantum yield, chlorophyll fluorescence parameters were measured weekly with a FluorCam FC800 (PSI, Bruno, Czech Republic) after a 20-minute dark adaptation. Maximum quantum yield (QY_max) was determined using the FluorCam 7 software.

Normality of data was checked using the Shapiro-Wilke’s test prior to checking homoskedasticity of variances using Levene’s Test in the R package ‘car’ (64). Growth rate (mm day^−1^) and total growth over the duration of the experiment were rank-transformed prior to two-way ANOVA to assess the influence of experimental temperature and donor microbiome. Fluorescence (Fv/Fm) was measured in moss as a proxy for photosynthetic activity throughout the experiment. However, to highlight the greatest differences in donor microbiome and experimental temperature combinations, only fluorescence data from the last week were used. Fluorescence data was also rank-transformed prior to using a two-way ANOVA. ⍰= 0.05 was used to denote statistical significance in both two-way ANOVA and Tukey’s HSD post-hoc analyses. Growth and fluorescence statistics were analyzed using R version 3.5.1 (65).

### 16s rDNA and ITS sequencing of community profiles

To characterize microbiomes of inocula and final microbiomes of the laboratory experiments, each sample was pulverized in liquid N_2_, and DNA was extracted from 50 mg of material using the DNeasy PowerPlant Pro Kit (Qiagen, Hilden, Germany). Extracted DNA was taken to the Genomics Core at the University of Tennessee, Knoxville, for library preparation and sequencing on an Illumina MiSeq. Libraries were prepped for the 16S rRNA gene using a two-step PCR approach with a mixture of custom 515F and 806R primers to characterize archaeal/bacterial communities, and for the *ITS2* gene region using a custom mixture of primers to characterize the fungal community (66). The initial PCR consisted of 2X KAPA HiFi HotStart Ready Mix Taq (Roche), 10 μM total for each forward and reverse primer combination, and approximately 50 ng of DNA. PCRs for 16S rRNA and ITS2 were performed separately. Reaction conditions were as follows: 3 minutes at 95°C; 25 cycles of 95°C for 30 seconds, 55°C for 30 seconds, and 72°C for 30 seconds; and a final extension at 72 degrees for 5 minutes. PCR products were purified using AMPure XP beads (Agencourt/Beckman Coulter, Indianapolis, IN, USA). Nextera XT indexes were ligated to the PCR products via a second, reduced-cycle PCR such that each sample had a unique combination of forward and reverse indexes, and the products were again purified using AMPure XP beads. Samples were pooled in equal concentrations and sequenced on the MiSeq along with negative control samples.

Microbial sequences were processed with QIIME 2 v 2018.11 platform (67). Paired sequences were demultiplexed with the plugin demux and quality-filtered (denoised, dereplicated, chimera-filtered, and pair-end merged) and processed into Sequence Variants (SVs) with the dada2 plugin (68). Taxonomy was assigned using a pre-trained Naive Bayes classifier based on Greengenes v13_8 (99% OTUs) that are trimmed to the 515F/806R primer pair for 16S rDNA and based on the UNITE (99% OTUs) for ITS2. Sequences assigned as chloroplast or mitochondria were removed. Microbial diversity was calculated based on a subsample of 19,000 sequences to fit the size of the smallest library. SV-based alpha diversity (Shannon diversity) and beta diversity (Bray-Curtis) were calculated using the phyloseq 1.30.0 (69) package in R (65). Beta diversity was visualized using nonmetric multidimensional scaling ordination (NMDS) based on Bray-Curtis similarity distances. A permutational multivariate analysis of variance (PERMANOVA) with 999 permutations was used to calculate the significance of clustering of samples by microbial and chamber treatment. Microbial diversity correlation with *Sphagnum* growth were assessed using Pearson correlation.

### Metagenomics of the starting inoculum

DNA for metagenomics was extracted using a modified CTAB method (70). The ambient- and warming-conditioned microbiome samples were sequenced as an Illumina TruSeq PCR-Free library on an Illumina 2500 in Rapid Run mode (paired-end, 2 × 150 nt). Raw sequences were processed using Atropos v. 1.1.17 (71) in Python v3.6.2 to remove adapter contamination and quality trimming. Quality trimming was performed at the Q20 level with read pairs, and if either read of a pair was < 75 bp after adapter removal and quality trimming, the read pair was discarded. Before metagenome assembly, we removed reads mapping to either the *S. angustifolium* or PhiX genomes using bbmap v38.22. The remaining metagenome reads for the ambient and warming samples were co-assembled using MEGAHIT v1.1.3 (72, 73) with default settings except --min-contig-len was set to 300. Trimmed paired-end reads were mapped to the MEGAHIT co-assembly with BamM v1.7.3 (http://ecogenomics.github.io/BamM/) (74). Putative genomes were binned from the co-assembled contigs using MetaBAT v2.12.1 (75) with – minContig set to 2000. Metagenome-assembled genomes (MAGs) were assessed for completeness, contamination, and strain contamination using checkM v1.0.12 (40). MAGs that were ≥70% complete with ≤5% contamination were kept for downstream analyses. To determine differences in the relative abundance of MAGs between ambient and warming metagenome samples, trimmed metagenomic reads were mapped to MAGs using BamM v.1.7.3 and normalized as counts per million mapped reads. Gene models were predicted for the co-assembled metagenomic contigs using Prodigal v2.6.3 (76) in anonymous mode, and MAGs were annotated using prokka v1.14.0 (77). Taxonomy was assigned to gene models using the lowest common ancestor algorithm implemented in DIAMOND v0.9.22 (41). Inferred amino acid sequences were searched against the NCBI non-redundant protein database (downloaded October 3, 2019) using default settings in DIAMOND BLASTp except as follows: --query-cover 85, --top 5, and --sensitive.

### Phylogenetic Analysis of Cyanobacterial MAGs

We downloaded 109 cyanobacterial RefSeq genomes from NCBI representing the currently sequenced diversity of clade A (*Oscillatoria/Arthrospira)* and clade B (*Nostoc/Anabaena/Cyanothece*), following the nomenclature of (43), and two outgroup taxa, *Acaryochloris* sp. CCMEE 5410 and *A. marina* MBIC11017. Additionally, we included six cyanobacterial isolates sequenced in (42), *Nostoc moss6, N. moss3, N. moss2, N. moss4, N. moss5,* and *N. sp. 996,* six MAGs from our metagenomic assembly labeled bin14, bin192, bin238, bin354, bin377, and bin384, and three binned genomes from the SPRUCE site, but outside the enclosures labeled bin109, bin297, and bin367. A concatenated alignment of inferred amino acids sequences from 31 proteins for the 125 cyanobacterial genomes was generated and trimmed using the AMPORA2 pipeline (78). Alignment sites containing only gaps and ambiguous characters were removed using FAST v1.6 (79). Molecular evolution model selection was performed with ModelFinder (80). Phylogenetic analysis was conducted with IQ-TREE v1.6.8 (81) using the cpREV+C60+F+R6 model. Node support was evaluated using 1000 SH-like approximate likelihood ratio test (82) replicates and 1000 UFboot2 replicates (81).

### Metatranscriptomics Profiling

Cryogenically stored samples from the end of the experiment were ground in liquid nitrogen, and total RNA was extracted using a method combining CTAB lysis buffer and the Spectrum Total Plant RNA extraction kit (Sigma, Darmstadt, Germany) as described previously (83). RNA quality and quantity were determined using a NanoDrop Spectrophotometer (Thermo Scientific, Waltham, MA, USA). Total RNA (3 μg) of three biological replicates was sent to Macrogen (Seoul, South Korea), where libraries were prepared and sequenced on an Illumina HiSeq.

Metatranscriptome reads were partitioned into *S. angustifolium* and microbial transcripts by mapping reads to the *S. angustifolium* v1.0 genome using bbmap v38.22. Microbial transcripts were processed using the SAMSA v2.2.0 pipeline (84), except that differential expressed SEED functional gene ontologies (85) were identified using limma-voom v3.11(86) with multiple testing correction using FDR. Taxonomic classification of microbial transcripts was performed by mapping reads to the metagenome assembly using BamM v1.7.3 and transferring the taxonomic classification of metagenomic gene models and MAG assignments to mapped transcripts.

To identify differentially expressed (DE) *S. angustifolium* genes, *S. angustifolium* read-pairs were mapped to S. *angustifolium* v1.0 reference genome using RSubread v2.3.0 (87) and analyzed using limma-voom v3.11. Enrichment of MapMan4 ontology bins (88) in the set of DE genes was determined using the MapMan desktop application v3.6.0RC1 (89). Statistical significance of MapMan ontology bins was determined using Kruskal–Wallis test with multiple testing correction using FDR in R v3.6.1. Log_2_(fold change) (LFC) of MapMan4 ontology bins were determined by averaging LFC across DE genes within each bin.

## Data Availability

Raw 16S and ITS sequence files can be found on NCBI using the BioProject ID PRJNA644113. Raw metagenome and metatranscriptome sequence files can be found on NCBI using the BioProject ID PRJNA644538.

## Acknowledgements

We are grateful for pre-submission comments from Dr. Gustaf Granath and field site maintenance from Robert Nettles III. Collection of starting microbial inocula was made possible through the SPRUCE project, which is supported by Office of Science; Biological and Environmental Research (BER); US Department of Energy (DOE), Grant/Award Number: DE-AC05–00OR22725. Experimentation, sample collection, and analyses were supported by the DOE BER Early Career Research Program. This research used resources of the Compute and Data Environment for Science (CADES) at the Oak Ridge National Laboratory. Oak Ridge National Laboratory is managed by UT-Battelle, LLC, for the US DOE under contract no. DE-AC05-00OR22725. AJS was supported by NSF DEB-1737899, 1928514. The work conducted by the US DOE Joint Genome Institute (JGI) is supported by the Office of Science of the US Department of Energy under Contract No. DE-AC02-05CH11231. We thank the DOE JGI and collaborators for pre-publication access to the *S. angustifolium* (formerly *S. fallax)* genome sequence.

## REFERENCES

1. van Breemen N, Nations U, Use I (1995) How Sphagnum bogs down other plants. Trends Ecol Evol (Personal Ed 10(7):270–275.

2. Clymo RS, Hayward PM (1982) The Ecology of Sphagnum. Bryophyte Ecology (Springer Netherlands), pp 229–289.

3. Yu Z, Loisel J, Brosseau DP, Beilman DW, Hunt SJ (2010) Global peatland dynamics since the Last Glacial Maximum. Geophys Res Lett 37(13):1–5.

4. Gorham E (1991) Northern peatlands: role in the carbon cycle and probable responses to climatic warming. Ecol Appl 1(2):182–195.

5. Dorrepaal E, et al. (2003) Summer warming and increased winter snow cover affect Sphagnum fuscum growth, structure and production in a sub-arctic bog. Glob Chang Biol 10(1):93–104.

6. Robroek BJM, Limpens J, Breeuwer A, Schouten MGC (2007) Effects of water level and temperature on performance of four Sphagnum mosses. Plant Ecol 190(1):97–107.

7. Gunnarsson U, Granberg G, Nilsson M (2004) Growth, production and interspecific competition in Sphagnum: effects of temperature, nitrogen and sulphur treatments on a boreal mire. New Phytol 163(2):349–359.

8. Robroek BJM, Limpens J, Breeuwer A, Crushell PH, Schouten MGC (2007) Interspecific competition between Sphagnum mosses at different water tables. Funct Ecol 21(4):805–812.

9. Bragazza L (2008) A climatic threshold triggers the die-off of peat mosses during an extreme heat wave. Glob Chang Biol 14(11):2688–2695.

10. Bragazza L, et al. (2016) Persistent high temperature and low precipitation reduce peat carbon accumulation. Glob Chang Biol 22(12):4114–4123.

11. Norby RJ, Childs J, Hanson PJ, Warren JM (2019) Rapid loss of an ecosystem engineer: Sphagnum decline in an experimentally warmed bog. Ecol Evol 9(September):12571–12585.

12. Kostka JE, et al. (2016) The Sphagnum microbiome: New insights from an ancient plant lineage. New Phytol 211(1):57–64.

13. Weston DJ, et al. (2014) Sphagnum physiology in the context of changing climate: emergent influences of genomics, modelling and host-microbiome interactions on understanding ecosystem function. Plant Cell Environ:n/a-n/a.

14. Lindo Z, Nilsson MC, Gundale MJ (2013) Bryophyte-cyanobacteria associations as regulators of the northern latitude carbon balance in response to global change. Glob Chang Biol 19(7):2022–2035.

15. Berg A, Danielsson Å, Svensson BH (2013) Transfer of fixed-N from N2-fixing cyanobacteria associated with the moss Sphagnum riparium results in enhanced growth of the moss. Plant Soil 362(1–2):271–278.

16. Basilier K (1979) Moss-Associated Nitrogen Fixation in Some Mire and Coniferous Forest Environments around Uppsala, Sweden. Lindbergia 5(2):84–88.

17. Basilier K, Granhall U, Stenström T-A (1978) Nitrogen Fixation in Wet Minerotrophic Moss Communities of a Subarctic Mire. Oikos 31(2):236.

18. Basilier K (1980) Fixation and Uptake of Nitrogen in Sphagnum Blue-Green Algal Associations. Oikos 34(2):239.

19. Granhall ULF, Hofsten A V (1976) Nitrogenase Activity in Relation to Intracellular Organisms in Sphagnum Mosses. Physiol Plant 36:88–94.

20. Vile M a., et al. (2014) N2-fixation by methanotrophs sustains carbon and nitrogen accumulation in pristine peatlands. Biogeochemistry. doi:10.1007/s10533-014-0019-6.

21. Larmoia T, et al. (2014) Methanotrophy induces nitrogen fixation during peatland development. Proc Natl Acad Sci U S A 111 (2):734–739.

22. Liebner S, Svenning MM (2013) Environmental transcription of mmoX by methane-oxidizing Proteobacteria in a subarctic palsa peatland. Appl Environ Microbiol 79(2):701–706.

23. Kip N, Winden J van, Pan Y (2010) Global prevalence of methane oxidation by symbiotic bacteria in peat-moss ecosystems. Nat Geosci 3(August):617–621.

24. Raghoebarsing AA, et al. (2005) Methanotrophic symbionts provide carbon for photosynthesis in peat bogs. Nature 436(7054):1153–1156.

25. Lamentowicz M, Mitchell EAD (2005) The ecology of testate amoebae (protists) in Sphagnum in north-western Poland in relation to peatland ecology. Microb Ecol 50(1):48–63.

26. Jassey VEJJ, et al. (2015) An unexpected role for mixotrophs in the response of peatland carbon cycling to climate warming. Sci Rep 5(November):16931.

27. Stough JMA, Kolton M, Kostka JE, Weston DJ, Pelletier DA (2018) crossm Diversity of Active Viral Infections within the Sphagnum Microbiome. 84(23):1–16.

28. Carrell AA, et al. (2019) Experimental warming alters the community composition, diversity, and N 2 fixation activity of peat moss (Sphagnum fallax) microbiomes. Glob Chang Biol (May):2993–3004.

29. Heath KD, Stock AJ, Stinchcombe JR (2010) Mutualism variation in the nodulation response to nitrate. J Evol Biol 23(11):2494–2500.

30. Baker DM, Freeman CJ, Wong JCY, Fogel ML, Knowlton N (2018) Climate change promotes parasitism in a coral symbiosis. ISME J 12(3):921–930.

31. Cunning R, Silverstein RN, Baker a. C (2015) Investigating the causes and consequences of symbiont shuffling in a multi-partner reef coral symbiosis under environmental change. Proc R Soc B Biol Sci 282(1809):20141725–20141725.

32. Bay LK, Doyle J, Logan M, Berkelmans R, Bay LK (2016) Recovery from bleaching is mediated by threshold densities of background thermo-tolerant symbiont types in a reef-building coral. R Soc open Sci 3(6):160322.

33. Howells EJ, Abrego D, Meyer E, Kirk NL, Burt JA (2016) Host adaptation and unexpected symbiont partners enable reef-building corals to tolerate extreme temperatures. Glob Chang Biol 22(8):2702–2714.

34. Rodriguez RJ, et al. (2008) Stress tolerance in plants via habitat-adapted symbiosis. ISME J 2(4):404–16.

35. Giauque H, Connor EW, Hawkes C V. (2019) Endophyte traits relevant to stress tolerance, resource use and habitat of origin predict effects on host plants. New Phytol 221(4):2239–2249.

36. Redman RS, et al. (2011) Increased fitness of rice plants to abiotic stress via habitat adapted symbiosis: A strategy for mitigating impacts of climate change. PLoS One 6(7):1–10.

37. Giauque H, Hawkes C V. (2013) Climate affects symbiotic fungal endophyte diversity and performance. Am J Bot 100(7):1435–1444.

38. Hanson PJ, et al. (2016) Intermediate-scale community-level flux of CO2 and CH4 in a Minnesota peatland: putting the SPRUCE project in a global context. Biogeochemistry 129(3):255–272.

39. Bowers RM, et al. (2017) Minimum information about a single amplified genome (MISAG) and a metagenome-assembled genome (MIMAG) of bacteria and archaea. Nat Biotechnol 35(8):725–731.

40. Parks DH, Imelfort M, Skennerton CT, Hugenholtz P, Tyson GW (2015) CheckM: Assessing the quality of microbial genomes recovered from isolates, single cells, and metagenomes. Genome Res 25(7):1043–1055.

41. Buchfink B, Xie C, Huson DH (2014) Fast and sensitive protein alignment using DIAMOND. Nat Methods 12(1):59–60.

42. Warshan D, et al. (2017) Feathermoss and epiphytic Nostoc cooperate differently: expanding the spectrum of plant–cyanobacteria symbiosis. ISME J:1–13.

43. Shih PM, et al. (2013) Improving the coverage of the cyanobacterial phylum using diversity-driven genome sequencing. Proc Natl Acad Sci U S A 110(3):1053–1058.

44. Cavicchioli R, et al. (2019) Scientists’ warning to humanity: microorganisms and climate change. Nat Rev Microbiol 17(September). doi:10.1038/s41579-019-0222-5.

45. Redman RS, Sheehan KB, Stout RG, Rodriguez RJ, Henson JM (2002) Thermotolerance Generated by Plant/Fungal Symbiosis. Science (80-) 298(5598):1581–1581.

46. Moya A, Ferrer M (2016) Functional Redundancy-Induced Stability of Gut Microbiota Subjected to Disturbance. Trends Microbiol 24(5):402–413.

47. Burke C, Steinberg P, Rusch D, Kjelleberg S, Thomas T (2011) Bacterial community assembly based on functional genes rather than species. Proc Natl Acad Sci U S A 108(34):14288–14293.

48. Apprill A (2020) The Role of Symbioses in the Adaptation and Stress Responses of Marine Organisms. Ann Rev Mar Sci 12(1):291–314.

49. Ziegler M, Seneca FO, Yum LK, Palumbi SR, Voolstra CR (2017) Bacterial community dynamics are linked to patterns of coral heat tolerance. Nat Commun 8:1–8.

50. Larkindale J, Mishkind M, Vierling E (2007) Plant Responses to High Temperature. Plant Abiotic Stress:100–144.

51. Mclellan CA, et al. (2007) A Rhizosphere Fungus Enhances Arabidopsis Thermotolerance through Production of an HSP90 Inhibitor 1. doi:10.1104/pp.107.101808.

52. Clarke SM, et al. (2009) Jasmonates act with salicylic acid to confer basal thermotolerance in Arabidopsis thaliana. New Phytol 182(1):175–187.

53. Yamamoto Y, et al. (2015) Functional analysis of allene oxide cyclase, MpAOC, in the liverwort Marchantia polymorpha. Phytochemistry 116(1):48–56.

54. Huang J, et al. (2010) Functional analysis of the Arabidopsis PAL gene family in plant growth, development, and response to environmental stress. Plant Physiol 153(4):1526–1538.

55. Kotak S, et al. (2007) Complexity of the heat stress response in plants. Curr Opin Plant Biol 10(3):310–316.

56. Conrath U, et al. (2006) Priming: Getting Ready for Battle. Mol Plant-Microbe Interact MPMI 19(10):1062–1071.

57. Ali SZ, et al. Pseudomonas sp. strain AKM-P6 enhances tolerance of sorghum seedlings to elevated temperatures. doi:10.1007/s00374-009-0404-9.

58. Meena KK, et al. (2017) Abiotic stress responses and microbe-mediated mitigation in plants: The omics strategies. Front Plant Sci 8:172.

59. Lau JA, Lennon JT (2012) Rapid responses of soil microorganisms improve plant fitness in novel environments. Proc Natl Acad Sci U S A 109(35):14058–62.

60. Weston DJ, et al. (2017) The Sphagnome Projects□: enabling ecological and evolutionary insights through a genus-level sequencing project Author Affiliations: 1–29.

61. Keuper F, et al. (2020) Carbon loss from northern circumpolar permafrost soils amplified by rhizosphere priming. Nat Geosci 13(8):560–565.

62. Gavazov K, et al. (2018) Vascular plant-mediated controls on atmospheric carbon assimilation and peat carbon decomposition under climate change. Glob Chang Biol 24(9):3911–3921.

63. Schneider CA, Rasband WS, Eliceiri KW (2012) NIH Image to ImageJ: 25 years of image analysis. Nat Methods 9(7):671–675.

64. Fox J, Weisberg S (2019) An R companion to applied regression (Third) (Sage, Thousand Oaks CA).

65. R Core Team (2015) R: A language and environment for statistical computing. Available at: http://www.r-project.org/.

66. Cregger MA, et al. (2018) The Populus holobiont: Dissecting the effects of plant niches and genotype on the microbiome. Microbiome 6(1):1–14.

67. Bolyen E, et al. (2019) Reproducible, interactive, scalable and extensible microbiome data science using QIIME 2. Nat Biotechnol 37(8):852–857.

68. Callahan BJ, et al. (2016) DADA2: High-resolution sample inference from Illumina amplicon data. Nat Methods 13(7):581–583.

69. McMurdie PJ, Holmes S (2013) phyloseq: An R Package for Reproducible Interactive Analysis and Graphics of Microbiome Census Data. PLoS One 8(4):e61217.

70. Carrell AA, Frank AC (2014) Pinus flexilis and Picea engelmannii share a simple and consistent needle endophyte microbiota with a potential role in nitrogen fixation. Front Microbiol 5(JULY):1–11.

71. Didion JP, Martin M, Collins FS (2017) Atropos: Specific, sensitive, and speedy trimming of sequencing reads. PeerJ 2017(8):1–19.

72. Li D, et al. (2016) MEGAHIT v1.0: A fast and scalable metagenome assembler driven by advanced methodologies and community practices. Methods 102:3–11.

73. Li D, Liu CM, Luo R, Sadakane K, Lam TW (2015) MEGAHIT: An ultra-fast single-node solution for large and complex metagenomics assembly via succinct de Bruijn graph. Bioinformatics 31(10):1674–1676.

74. Li H, Durbin R (2009) Fast and accurate short read alignment with Burrows-Wheeler transform. 25(14):1754–1760.

75. Kang DD, et al. (2019) MetaBAT 2: an adaptive binning algorithm for robust and efficient genome reconstruction from metagenome assemblies. PeerJ 7:e7359.

76. Hyatt D, et al. (2010) Prodigal: prokaryotic gene recognition and translation initiation site identification Available at: http://www.biomedcentral.com/1471-2105/11/119 [Accessed July 6, 2020].

77. Seemann T (2014) Genome analysis Prokka: rapid prokaryotic genome annotation. 30(14):2068–2069.

78. Wu M, Scott AJ (2012) Phylogenomic analysis of bacterial and archaeal sequences with AMPHORA2. Bioinforma Appl NOTE 28(7):1033–1034.

79. Lawrence TJ, et al. (2015) FAST: FAST Analysis of Sequences Toolbox. Front Genet 6(MAY):172.

80. Kalyaanamoorthy S, Minh BQ, Wong TKF, Von Haeseler A, Jermiin LS (2017) ModelFinder: Fast model selection for accurate phylogenetic estimates. Nat Methods 14(6):587–589.

81. Quang B, Anh M, Nguyen T, Von Haeseler A Ultrafast Approximation for Phylogenetic Bootstrap. doi:10.1093/molbev/mst024.

82. St’ S, et al. (2010) New Algorithms and Methods to Estimate Maximum-Likelihood Phylogenies: Assessing the Performance of PhyML 3.0. Syst Biol 59(3):307–321.

83. Timm CM, et al. (2016) Two poplar-associated bacterial isolates induce additive favorable responses in a constructed plant-microbiome system. Front Plant Sci 7(APR2016):1–10.

84. Westreich ST, Korf I, Mills DA, Lemay DG (2016) SAMSA: a comprehensive metatranscriptome analysis pipeline. doi:10.1186/s12859-016-1270-8.

85. Overbeek R, et al. The SEED and the Rapid Annotation of microbial genomes using Subsystems Technology (RAST). doi:10.1093/nar/gkt1226.

86. Ritchie ME, et al. (2015) limma powers differential expression analyses for RNA-sequencing and microarray studies. Nucleic Acids Res 43(7). doi:10.1093/nar/gkv007.

87. Liao Y, Smyth GK, Shi W (2019) The R package Rsubread is easier, faster, cheaper and better for alignment and quantification of RNA sequencing reads. Nucleic Acids Res 47(8). doi:10.1093/nar/gkz114.

88. Schwacke R, et al. (2019) MapMan4: A Refined Protein Classification and Annotation Framework Applicable to Multi-Omics Data Analysis. Mol Plant 12:879–892.

89. Thimm O, et al. (2004) mapman: a user-driven tool to display genomics data sets onto diagrams of metabolic pathways and other biological processes. Plant J 37(6):914–939.

