## Supplementary material for "*Sphagnum* peat moss thermotolerance is modulated by the microbiome": SI Appendix

### Environmental symbiosis drives plant resilience to warming

#### This PDF file includes:

Figs. S1 to S7
Tables S1 to S13
Legend for Dataset S1

#### Other supplementary materials for this manuscript include the following:

Dataset S1

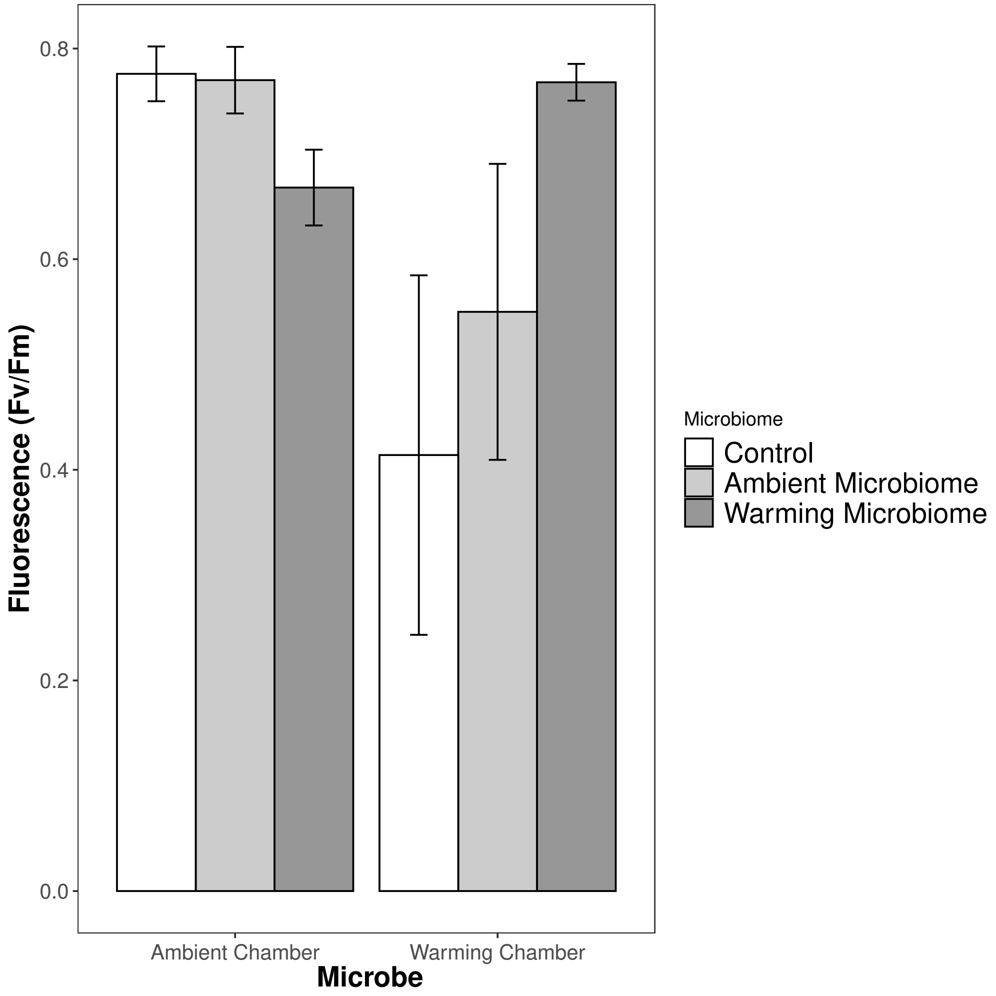


**Fig. S1.** Average moss fluorescence in 2016 at the end of the experiment with error bars representing standard error of the mean.

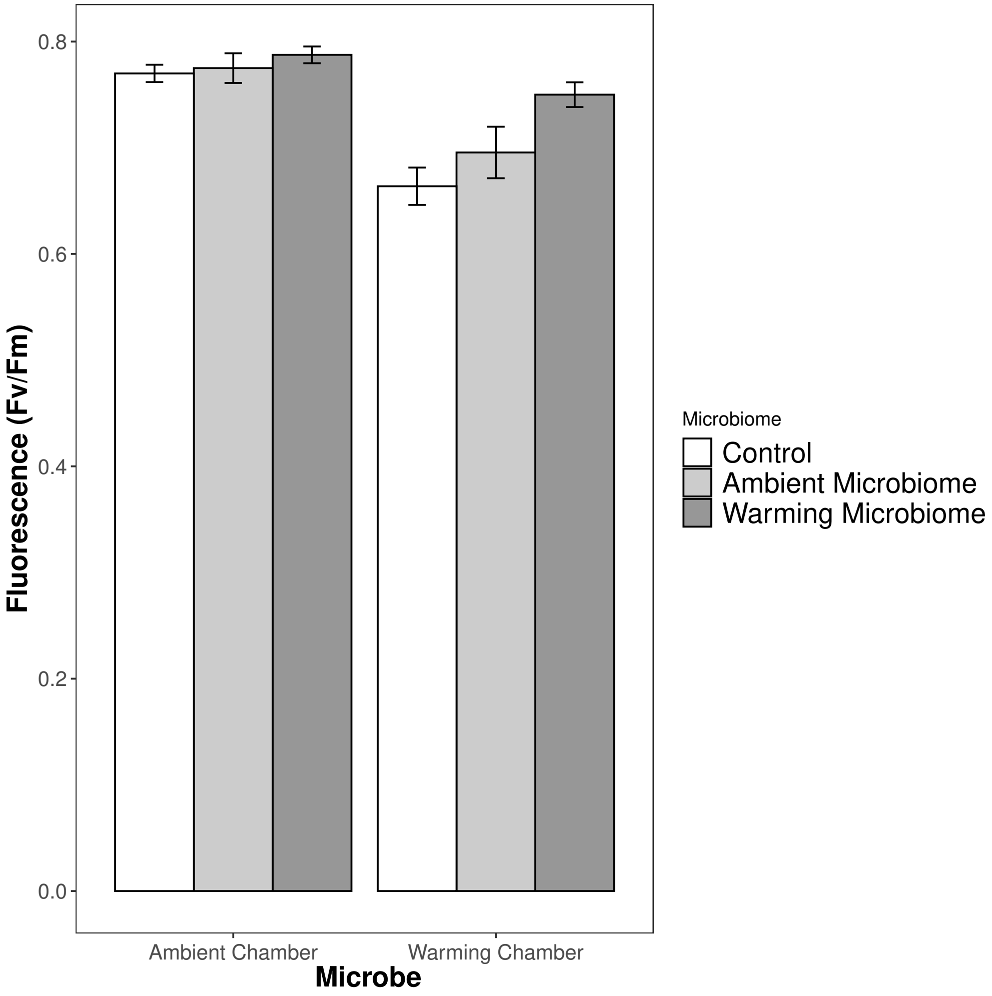


**Fig. S2.** Average moss fluorescence in 2017 at the end of the experiment with error bars representing standard error of the mean.

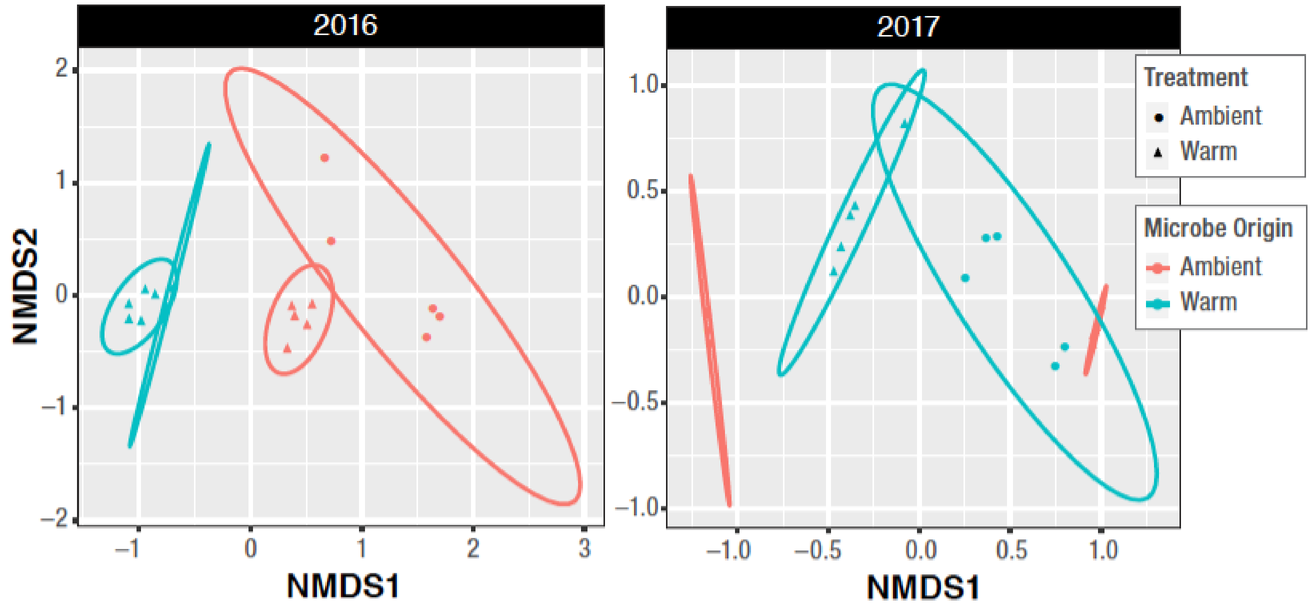


**Fig. S3.** Non-metric multidimensional scaling (NMDS) ordination of the microbiome Bray-Curtis distance matrix. Points that are closer together on the ordination have communities that are more similar. Each point corresponds to a sample, color corresponds to the microbiome thermal origin from ambient and warm field plots. Shapes correspond to the chamber that axenic plants were incubated at ambient or warm temperatures.

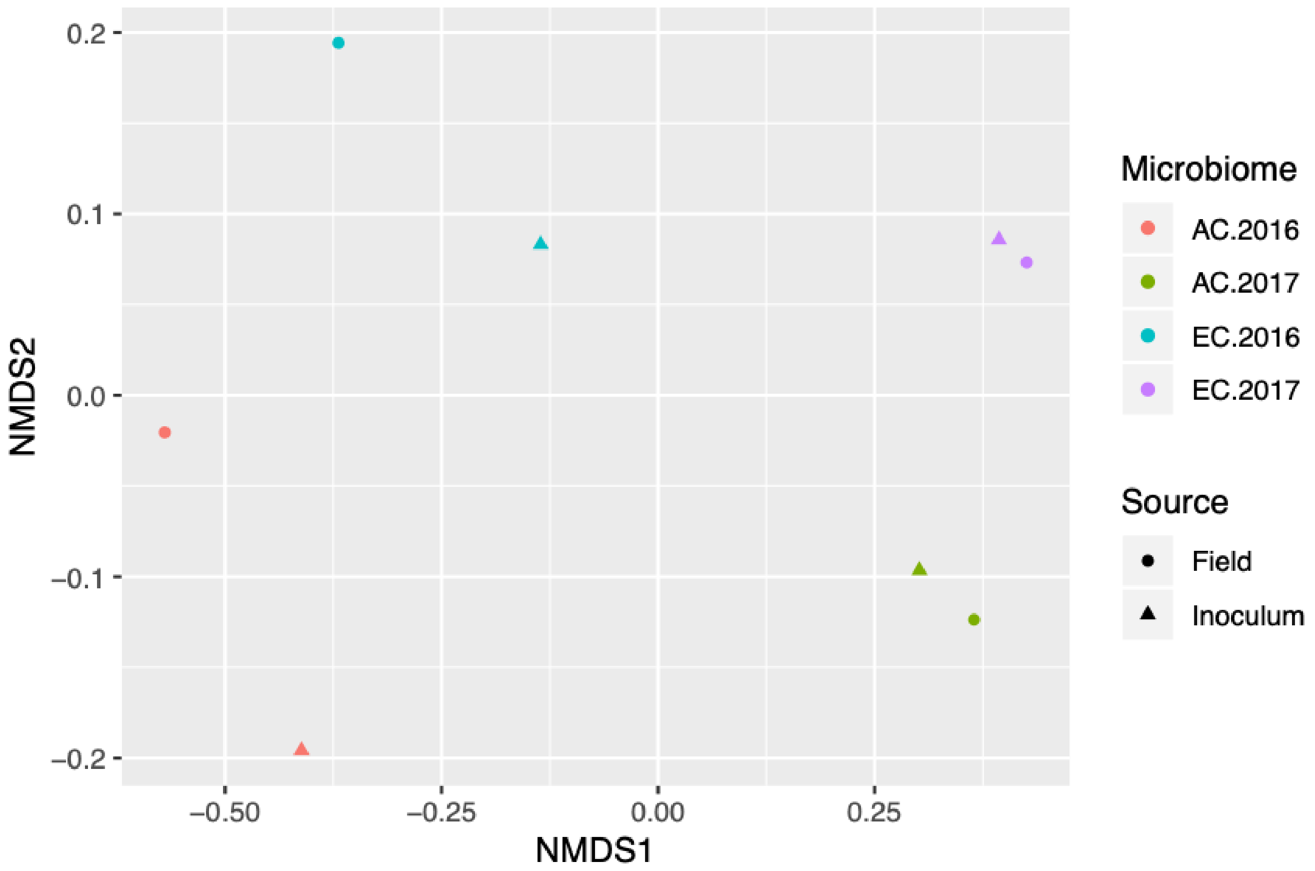


**Fig. S4.** Non-metric multidimensional scaling (NMDS) ordination of the Bray-Curtis distance matrix of SSU rRNA gene sequences rarefied to 19000 reads per sample. Points that are closer together on the ordination have communities that are more similar. Each point corresponds to a sample, color corresponds to the ambient (AC) and elevated (EC) SPRUCE enclosures and the year of collection, and shapes correspond to Sphagnum microbiomes from the SPRUCE site directly from field samples (field) or after the microbiomes were isolated from the field samples to be used as starting inoculum for incubation experiments in the laboratory.

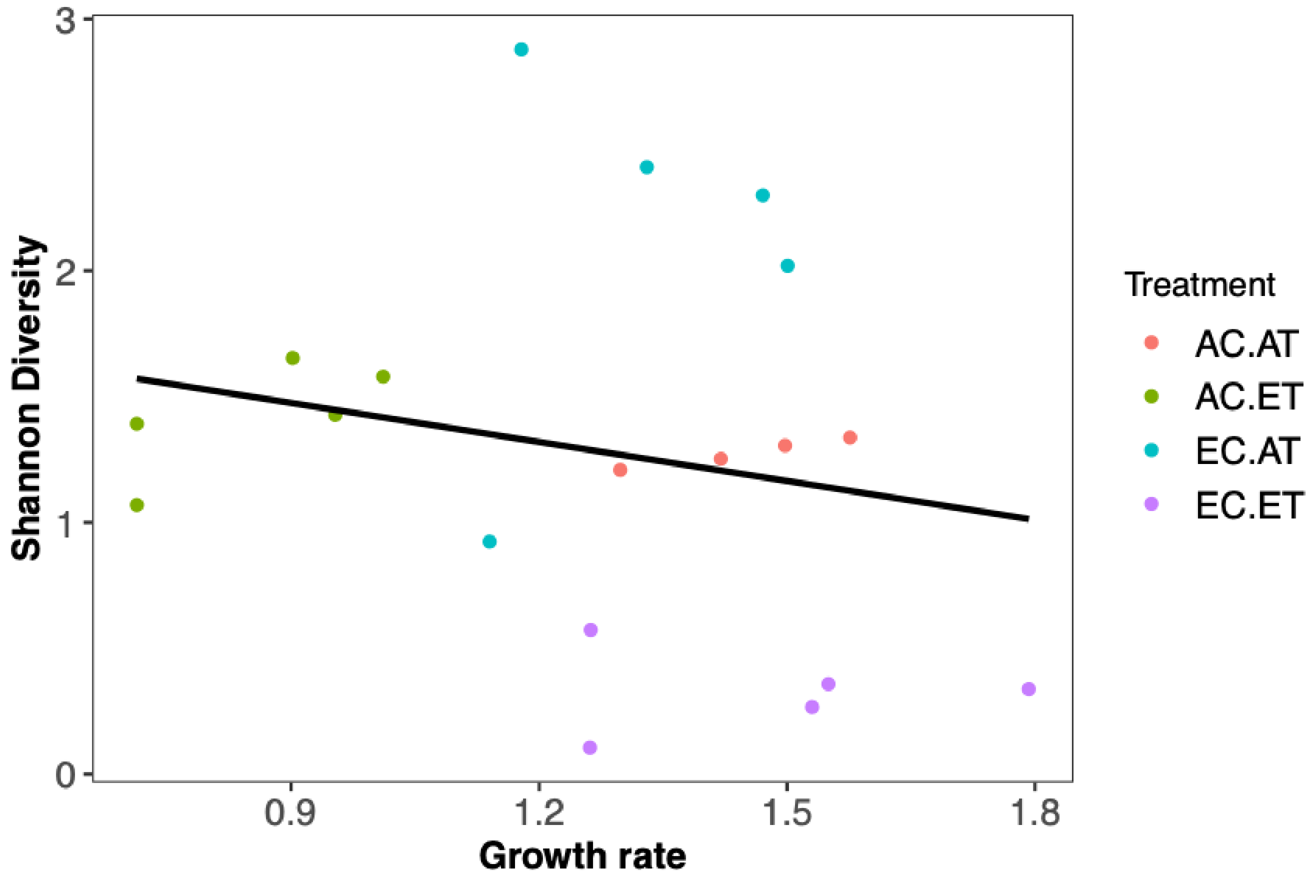


**Fig. S5.** Linear correlation of plant growth rate and fungal (ITS) Shannon diversity at the conclusion of the experiment. Each point corresponds to a sample with colors indicating ambient chamber/ambient microbiome (AC.AT), ambient chamber/warming microbiome (EC.AT), warming chamber/ambient microbiome (AC.ET), or warming chamber/warming microbiome (EC.ET).

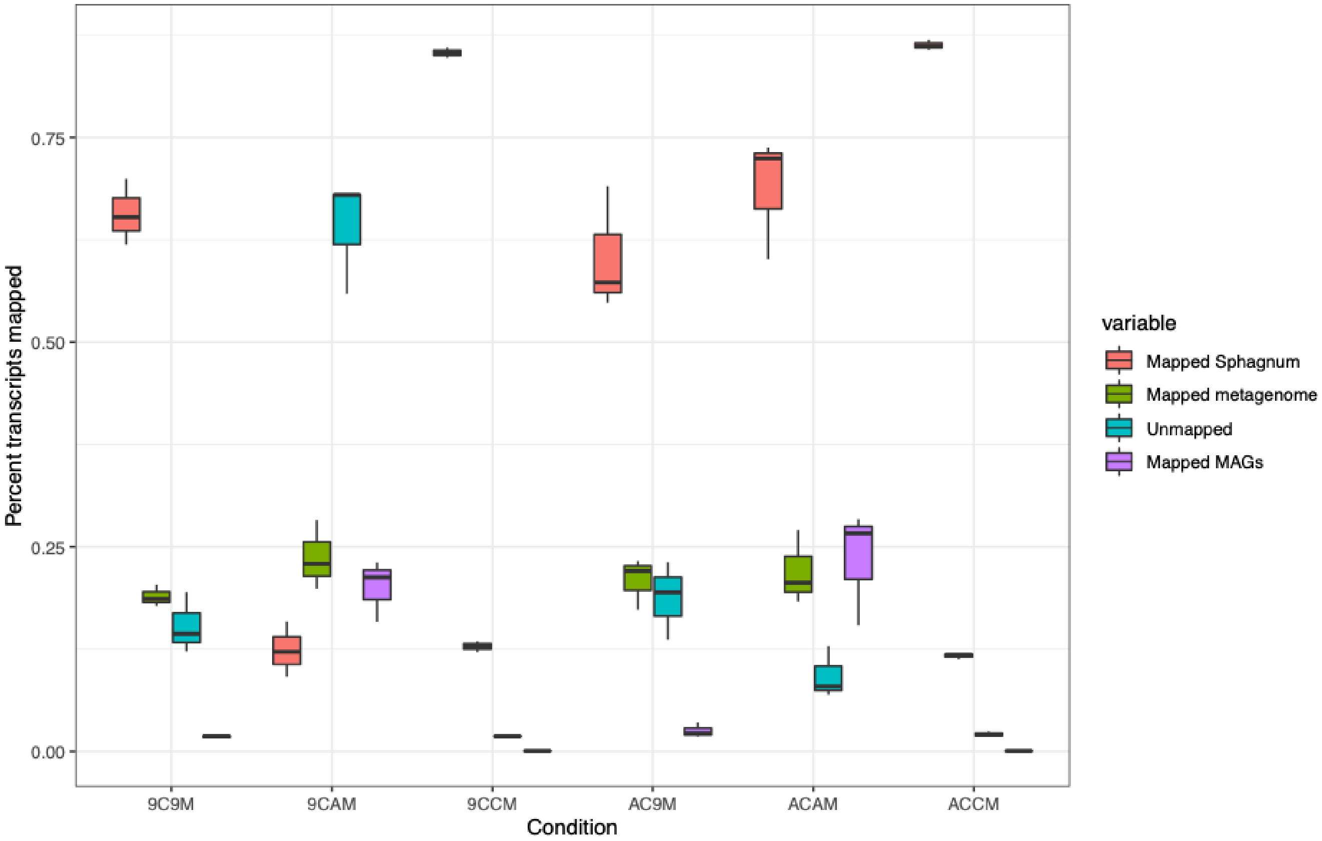


**Fig. S6.** Boxplot of percent of metatranscriptomic reads mapping to Sphagnum fallax (red), metagenome assembly, and metagenome assembled genomes (purple), or not mapping (blue) for ambient chamber/ambient microbiome (ACAM), ambient chamber/warming microbiome (AC9M), warming chamber/ambient microbiome (9CAM), warming chamber/warming microbiome (9C9M), ambient chamber/no microbiome addition (ACCM), and warming chamber/no microbiome addition (9CCM). Boxes span from the first to the third quartile, black bar indicates median, and whiskers span 1.5 * interquartile range.

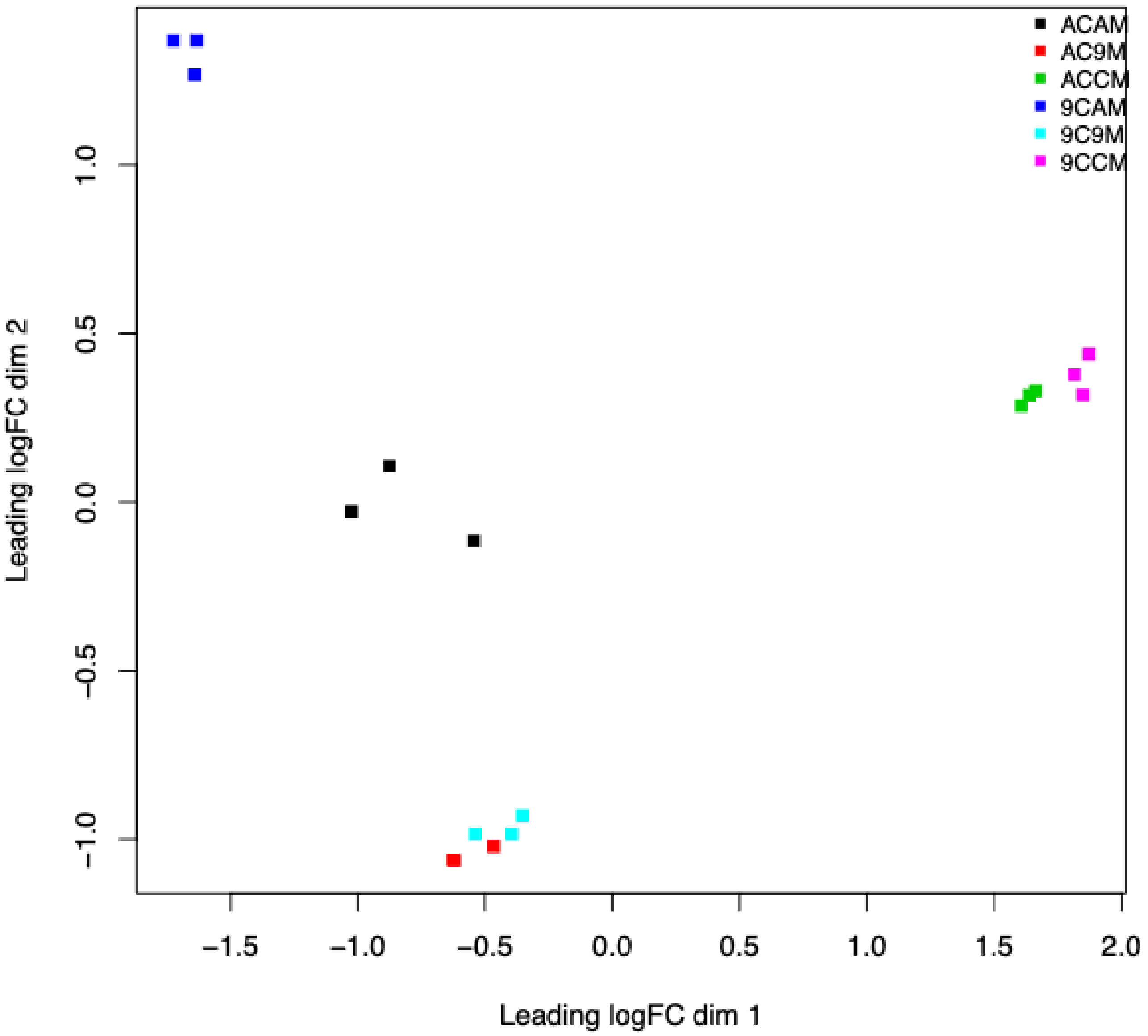


**Fig. S7.** MDS based on the top 500 most variable SEED level 3 categories of the microbial partition of the metatranscriptome. Abbreviations are defined in Fig. S6

**Table S1. Incubation temperature and light cycle for 2016 and 2017 laboratory experiments. Temperatures were determined from June 2016 average temperature for 6 hour blocks at the SPRUCE site.**

| Treatment Chamber | Time | Temperature (C) | Light |
|---|---|---|---|
| Ambient +0C | 0:00 | 13 | No |
| | 6:00 | 18 | Yes |
| | 12:00 | 21 | Yes |
| | 18:00 | 15 | No |
| Ambient +9C | 0:00 | 22 | No |
| | 6:00 | 27 | Yes |
| | 12:00 | 30 | Yes |
| | 18:00 | 24 | No |

**Table S2. Summary growth rate and total moss growth over 4 weeks.**

| Year | Chamber | Microbiome origin | N | Mean growth rate (mm/day) | SD growth rate | SE growth rate | Mean growth (mm) | SD growth | SE growth |
|---|---|---|---|---|---|---|---|---|---|
| 2016 | Ambient | Ambient | 5 | 1.664 | 0.448 | 0.200 | 46.601 | 12.534 | 5.605 |
| | Ambient | Control | 5 | 1.060 | 0.718 | 0.321 | 29.687 | 20.104 | 8.991 |
| | Ambient | Warming | 5 | 1.098 | 0.786 | 0.352 | 30.736 | 22.017 | 9.846 |
| | Warming | Ambient | 5 | 0.887 | 0.569 | 0.255 | 24.849 | 15.936 | 7.127 |
| | Warming | Control | 5 | 0.202 | 0.202 | 0.090 | 5.645 | 5.645 | 2.525 |
| | Warming | Warming | 5 | 1.759 | 0.418 | 0.187 | 49.251 | 11.692 | 5.229 |
| 2017 | Ambient | Ambient | 12 | 1.407 | 0.288 | 0.083 | 29.548 | 6.057 | 1.749 |
| | Ambient | Control | 12 | 1.088 | 0.117 | 0.034 | 22.844 | 2.455 | 0.709 |
| | Ambient | Warming | 12 | 1.311 | 0.186 | 0.054 | 27.537 | 3.906 | 1.128 |
| | Warming | Ambient | 12 | 0.593 | 0.455 | 0.131 | 12.455 | 9.564 | 2.761 |
| | Warming | Control | 12 | 0.188 | 0.179 | 0.052 | 3.948 | 3.758 | 1.085 |
| | Warming | Warming | 12 | 1.460 | 0.170 | 0.049 | 30.662 | 3.563 | 1.028 |

**Table S3. Two-way ANOVA tables of total moss growth. Data were ranked transformed prior to analysis.**

| | Sum of Squares | DF | F-value | P-value |
|---|---|---|---|---|
| 2016 | | | | |
| Intercept | 2420.00 | 1 | 53.226 | <0.05 |
| Chamber | 211.60 | 1 | 4.654 | <0.05 |
| Microbe | 170.53 | 2 | 1.875 | 0.175 |
| Chamber:Microbiome | 516.47 | 2 | 5.680 | <0.05 |
| Residuals | 1091.20 | 24 | | |
| 2017 | | | | |
| Intercept | 32865 | 1 | 308.448 | <0.05 |
| Chamber | 7633 | 1 | 71.634 | <0.05 |
| Microbe | 2477 | 2 | 11.621 | <0.05 |
| Chamber:Microbiome | 6823 | 2 | 32.016 | <0.05 |
| Residuals | 7032 | 66 | | |

**Table S4. Percent change of total moss growth between microbiome transfers relative to moss without a microbiome within the same chamber calculated as: $(\text{no microbiome} - \text{treatment microbiome})/\text{no microbiome}$.**

| Year | Chamber | Microbiome | Percent change | p-value |
|---|---|---|---|---|
| 2016 | Ambient | Ambient | 36% | ns |
| | Ambient | Warming | 3% | ns |
| | Warming | Ambient | 77% | ns |
| | Warming | Warming | 89% | <0.05 |
| 2017 | Ambient | Ambient | 23% | <0.05 |
| | Ambient | Warming | 17% | <0.05 |
| | Warming | Ambient | 68% | ns |
| | Warming | Warming | 87% | <0.05 |

**Table S5. Two-way ANOVA tables of moss fluorescence (Fv/Fm). Data were ranked transformed prior to analysis.**

| | Sum of squares | DF | F-value | P-value |
|---|---|---|---|---|
| 2016 | | | | |
| Intercept | 2000.00 | 1 | 37.433 | <0.05 |
| Chamber | 225.62 | 1 | 4.223 | 0.051 |
| Microbe | 352.43 | 2 | 3.298 | 0.054 |
| Chamber:Microbiome | 822.02 | 2 | 7.693 | <0.05 |
| Residuals | 1282.30 | 24 | | |
| 2017 | | | | |
| Intercept | 17328.0 | 1 | 40.135 | <0.05 |
| Chamber | 222.0 | 1 | 0.514 | 0.476 |
| Microbe-Origin | 577.8 | 2 | 0.669 | 0.516 |
| Chamber:Microbiome | 328.8 | 2 | 0.381 | 0.685 |
| Residuals | 28494.8 | 66 | | |

**Table S6. Percent change of fluroesence at harvest between microbiome transfers relative to moss without a microbiome within the same chamber calculated as: $(\text{no microbiome} - \text{treatment microbiome})/\text{no microbiome}$.**

| Year | Chamber | Microbiome | Percent change | p-value |
|---|---|---|---|---|
| 2016 | Ambient | Ambient | -0.78% | ns |
| | Ambient | Warming | -16.17% | ns |
| | Warming | Ambient | 24.73% | ns |
| | Warming | Warming | 46.09% | ns |
| 2017 | Ambient | Ambient | 0.65% | ns |
| | Ambient | Warming | 2.28% | ns |
| | Warming | Ambient | 4.59% | ns |
| | Warming | Warming | 11.47% | ns |

**Table S7. Community abundance table for starting inoculums isolated from the ambient (AC) and warming (EC) SPRUCE enclosures and the year of collection.**

| Class | AC.2016 | AC.2017 | EC.2016 | EC.2017 |
|---|---|---|---|---|
| c__[Fimbriimonadia] | NA | 0.0048 | NA | 0.0044 |
| c__[Methylacidiphilae] | 0.0067 | 0.0422 | NA | 0.0175 |
| c__[Pedosphaerae] | 0.017 | 0.0644 | 0.0059 | 0.0463 |
| c__[Saprospirae] | 0.0227 | 0.0447 | 0.0441 | 0.0329 |
| c__[Spartobacteria] | 0.0044 | 0.0303 | 0.0011 | 0.018 |
| c__4C0d-2 | 0.0015 | 0.0058 | NA | 0.0026 |
| c__Acidimicrobiia | 0.0049 | 0.0065 | NA | 0.0066 |
| c__Acidobacteriia | 0.124 | 0.1127 | 0.0641 | 0.1329 |
| c__Actinobacteria | 0.0168 | 0.0093 | 0.0039 | 0.0035 |
| c__Alphaproteobacteria | 0.3211 | 0.2954 | 0.0805 | 0.3033 |
| c__Armatimonadia | NA | 0.0031 | NA | 0.0033 |
| c__Bacilli | NA | NA | 0.0151 | NA |
| c__Bacteroidia | 0.008 | NA | 0.0016 | NA |
| c__Betaproteobacteria | 0.0642 | 0.0562 | 0.1706 | 0.0803 |
| c__Chlamydiia | 0.0141 | 0.0082 | 0.0047 | 0.0107 |
| c__Chthonomonadetes | NA | 0.0031 | NA | 0.0061 |
| c__Clostridia | 0.1609 | NA | 0.0654 | NA |
| c__Coriobacteriia | 0.0018 | NA | NA | NA |
| c__Cytophagia | NA | 0.0018 | NA | 0.0047 |
| c__Deltaproteobacteria | 0.0057 | 0.0272 | 0.0014 | 0.0192 |
| c__Elusimicrobia | 0.0018 | 0.0014 | NA | 0.0015 |
| c__Flavobacteriia | NA | NA | 0.0013 | 0.0014 |
| c__Gammaproteobacteria | 0.103 | 0.1115 | 0.4248 | 0.1448 |
| c__Holophagae | NA | NA | 0.0259 | NA |
| c__Methanobacteria | 0.0421 | NA | 0.0088 | NA |
| c__Methanomicrobia | 0.0014 | NA | NA | NA |
| c__Nostocophycideae | 0.0053 | 0.0189 | NA | 0.0173 |
| c__Opitutae | 0.0111 | 0.0127 | 0.0019 | 0.0087 |
| c__Phycisphaerae | 0.0037 | 0.0232 | NA | 0.015 |
| c__Planctomycetia | 0.0076 | 0.02 | NA | 0.0233 |
| c__SBRH58 | NA | 0.001 | NA | NA |
| c__SJA-4 | 0.0011 | 0.0022 | NA | 0.0011 |
| c__Solibacteres | 0.0109 | 0.0237 | NA | 0.047 |
| c__Sphingobacteriia | 0.0267 | 0.0531 | 0.0722 | 0.0377 |
| c__Thermoleophilia | 0.0025 | 0.0048 | NA | 0.0027 |
| c__Thermoplasmata | 0.0019 | NA | NA | NA |
| c__TM7-1 | NA | 0.0016 | NA | NA |
| c__vadinHA49 | NA | 0.0017 | NA | NA |
| c__Verrucomicrobiae | NA | 0.002 | NA | 0.0013 |
| c__ZB2 | NA | 0.0011 | NA | NA |
| Other | 0.007 | 0.0054 | 0.0066 | 0.0062 |

**Table S8. Community abundance table for samples at the end of the experiment for the ambient chamber/ambient microbiome (AC.AT), ambient chamber/warming microbiome (EC.AT), warming chamber/ambient microbiome (AC.ET), and warming chamber/warming microbiome (EC.ET).**

| Year | 2016 | | | | 2017 | | | |
|---|---|---|---|---|---|---|---|---|
| Class | AC.AT | AC.ET | EC.AT | EC.ET | AC.AT | AC.ET | EC.AT | EC.ET |
| c__[Methylacidiphilae] | NA | NA | 0.024 | NA | 0.028 | NA | NA | NA |
| c__[Pedosphaerae] | 0.083 | 0.025 | 0.032 | 0.024 | 0.043 | NA | NA | NA |
| c__[Saprospirae] | 0.024 | 0.094 | 0.083 | NA | 0.071 | 0.261 | 0.316 | 0.19 |
| c__[Spartobacteria] | 0.026 | 0.211 | 0.086 | NA | 0.053 | NA | NA | 0.026 |
| c__Acidobacteriia | 0.089 | 0.066 | 0.104 | 0.055 | 0.074 | NA | NA | NA |
| c__Actinobacteria | NA | NA | NA | NA | 0.026 | NA | NA | NA |
| c__Alphaproteobacteria | 0.288 | 0.189 | 0.185 | 0.221 | 0.256 | 0.146 | 0.159 | 0.258 |
| c__Betaproteobacteria | 0.111 | 0.202 | 0.229 | 0.339 | 0.105 | 0.049 | 0.153 | 0.116 |
| c__Clostridia | 0.089 | NA | NA | 0.119 | NA | NA | NA | NA |
| c__Deltaproteobacteria | NA | NA | NA | NA | 0.021 | NA | 0.036 | 0.021 |
| c__Flavobacteriia | NA | NA | NA | NA | NA | 0.07 | 0.074 | NA |
| c__Gammaproteobacteria | 0.091 | 0.046 | 0.059 | 0.052 | 0.071 | NA | 0.034 | 0.097 |
| c__Methanobacteria | 0.023 | NA | NA | 0.03 | NA | NA | NA | NA |
| c__Nostocophycideae | NA | NA | NA | NA | 0.136 | NA | NA | NA |
| c__Opitutae | 0.027 | NA | NA | NA | NA | NA | NA | 0.054 |
| c__SJA-4 | NA | NA | NA | NA | 0.024 | NA | NA | NA |
| c__Solibacteres | NA | 0.029 | 0.021 | NA | NA | NA | NA | 0.063 |
| c__Sphingobacteriia | NA | NA | 0.036 | 0.03 | 0.029 | NA | NA | 0.069 |
| c__Synechococcophycideae | NA | NA | NA | NA | NA | 0.419 | 0.133 | NA |
| c__Verrucomicrobiae | NA | 0.04 | 0.057 | NA | NA | NA | 0.026 | NA |
| Other | 0.149 | 0.097 | 0.084 | 0.13 | 0.064 | 0.056 | 0.07 | 0.107 |

**Table S9. Fungal (ITS) phylum level taxonomy of microbiomes at the end of the laboratory experiment. Column headings are defined in table S8.**

| Phylum | AC.AT | AC.ET | EC.AT | EC.ET |
|---|---|---|---|---|
| Ascomycota | 1.00 | 1.00 | 0.99 | 1.00 |
| Other | 0.00 | 0.00 | 0.01 | 0.00 |

**Table S10. Fungal (ITS) class level taxonomy of microbiomes at the end of the laboratory experiment. Column headings are defined in table S8.**

| Class | AC.AT | AC.ET | EC.AT | EC.ET |
|---|---|---|---|---|
| Dothideomycetes | 0.06 | 0.03 | 0.31 | NA |
| Leotiomycetes | 0.93 | 0.26 | 0.47 | 0.99 |
| Sordariomycetes | NA | 0.71 | 0.20 | NA |
| Other | 0.01 | 0.00 | 0.03 | 0.01 |

**Table S11. Descriptive statistics and log2 fold change of metagenome assembled genomes (MAG). Marker lineage, completeness, contamination, and strain heterogeneity were calculated using checkM. Log2 fold change between warming and ambient metagenomic samples were determined by mapping reads metagenomic reads back to MAGs using BAMM and calculating counts per million mapped reads.**

| Bin # | Marker lineage | Completeness | Contamination | Strain heterogeneity | log2FC (warming/ambient) |
|---|---|---|---|---|---|
| bin.354 | Cyanobacteria | 98.22 | 0.00 | 0.00 | 13.1807 |
| bin.259 | Chlamydiales | 97.97 | 0.68 | 0.00 | -11.6103 |
| bin.156 | Bacteroidetes | 94.77 | 0.27 | 66.67 | -10.5337 |
| bin.104 | Bacteroidetes | 92.42 | 2.10 | 10.00 | -8.5054 |
| bin.192 | Cyanobacteria | 88.92 | 0.84 | 0.00 | 8.1474 |
| bin.99 | Burkholderiales | 98.30 | 2.10 | 18.18 | 7.1546 |
| bin.287 | Chlamydiae | 88.68 | 0.68 | 0.00 | 6.4135 |
| bin.23 | Acidobacteriaceae | 93.41 | 3.42 | 28.57 | -5.6312 |
| bin.15 | Bacteroidetes | 93.79 | 2.94 | 33.33 | 5.6232 |
| bin.184 | Acidobacteriaceae | 83.10 | 1.71 | 50.00 | 5.5522 |
| bin.56 | Acidobacteriaceae | 91.64 | 4.00 | 42.86 | -5.4472 |
| bin.309 | Burkholderiales | 96.81 | 2.12 | 55.56 | -5.1306 |
| bin.47 | Acidobacteriaceae | 90.80 | 3.45 | 20.00 | -5.1036 |
| bin.81 | Acidobacteriaceae | 73.28 | 2.59 | 100.00 | -5.0201 |
| bin.292 | Burkholderiales | 75.00 | 1.72 | 100.00 | -4.9334 |
| bin.17 | Rhodospirillales | 94.96 | 1.62 | 50.00 | -4.5374 |
| bin.121 | Alphaproteobacteria | 94.64 | 5.00 | 13.64 | -4.2686 |
| bin.167 | Bradyrhizobiaceae | 93.76 | 3.89 | 40.91 | -3.5383 |
| bin.313 | Burkholderiales | 78.47 | 2.66 | 38.46 | 3.3494 |
| bin.59 | Chlamydiae | 94.59 | 0.68 | 0.00 | -3.0997 |
| bin.298 | Chlamydiae | 70.78 | 1.91 | 0.00 | 2.9728 |
| bin.229 | Rhodospirillales | 92.89 | 3.69 | 52.94 | -2.9154 |
| bin.72 | Bacteroidetes | 77.35 | 1.48 | 0.00 | 2.8915 |
| bin.14 | Cyanobacteria | 99.00 | 0.63 | 0.00 | -2.8049 |
| bin.79 | Rhizobiales | 84.70 | 3.33 | 57.14 | 2.7164 |
| bin.221 | Rhizobiales | 97.44 | 4.23 | 70.59 | -2.6230 |
| bin.33 | Gammaproteobacteria | 89.23 | 4.73 | 38.10 | -2.5113 |
| bin.120 | Betaproteobacteria | 88.46 | 1.33 | 0.00 | -2.4386 |
| bin.208 | Rhodospirillales | 76.88 | 2.99 | 77.78 | 2.2918 |
| bin.326 | Verrucomicrobia | 99.27 | 4.26 | 40.00 | 2.1862 |
| bin.31 | Gammaproteobacteria | 80.17 | 0.00 | 0.00 | -2.0919 |
| bin.302 | Rhodospirillales | 98.64 | 4.34 | 71.43 | 1.4258 |
| bin.255 | Bacteroidetes | 86.59 | 2.73 | 11.11 | -0.9504 |
| bin.189 | Rhodospirillales | 95.32 | 3.24 | 45.45 | -0.8350 |
| bin.265 | Verrucomicrobia | 97.29 | 3.99 | 0.00 | -0.8016 |
| bin.286 | Alphaproteobacteria | 93.01 | 1.02 | 33.33 | 0.7232 |
| bin.247 | Gammaproteobacteria | 96.46 | 3.83 | 33.33 | -0.6157 |
| bin.243 | Verrucomicrobiae | 93.68 | 0.68 | 0.00 | 0.5466 |
| bin.202 | Rhodospirillales | 92.42 | 0.72 | 33.33 | 0.4583 |
| bin.289 | Candidate phylum TM6 | 83.15 | 1.12 | 100.00 | 0.4233 |
| bin.92 | Rhodospirillales | 99.10 | 1.41 | 20.00 | 0.2420 |
| bin.339 | Acidobacteriaceae | 88.70 | 2.78 | 42.86 | -0.1816 |
| bin.132 | Bacteroidetes | 93.04 | 3.25 | 55.56 | 0.1609 |
| bin.223 | Rhodospirillales | 92.83 | 4.81 | 7.69 | -0.0729 |
| bin.198 | Actinomycetales | 98.48 | 2.27 | 40.00 | 0.0341 |

**Table S12. Number of paired-end metatranscriptomic reads passing quality control. Quality trimming was performed at the Q20 level and if either read of a pair was <75 bp after adapter removal and quality trimming the read pair was discarded.**

| Chamber | Microbiome | Rep | Counts of paired-end reads passing QC |
|---|---|---|---|
| Ambient | Ambient | 1 | 49,339,226 |
| | | 2 | 35,987,608 |
| | | 3 | 39,865,568 |
| Ambient | Warming | 1 | 33,372,123 |
| | | 2 | 31,528,203 |
| | | 3 | 41,055,820 |
| Ambient | Control | 1 | 29,530,155 |
| | | 2 | 37,222,244 |
| | | 3 | 31,634,008 |
| Warming | Ambient | 1 | 56,084,411 |
| | | 2 | 48,673,151 |
| | | 3 | 42,032,880 |
| Warming | Warming | 1 | 46,801,800 |
| | | 2 | 35,071,333 |
| | | 3 | 41,731,001 |
| Warming | Control | 1 | 45,655,644 |
| | | 2 | 49,482,126 |
| | | 3 | 41,676,823 |

**Table S13. Log2 fold change of SEED subsystem level 3 gene ontologies for the microbial fraction of metatranscriptomic reads. Read mapping and feature counts were determined using the SAMSA pipeline and differential expression analysis was performed with limma-voom. Abbreviations are defined as ambient chamber/ambient temperature conditioned microbiome (ACAM), +9C chamber/ambient temperature conditioned microbiome (9CAM), Ambient chamber/+9C temperature conditioned microbiome (AC9M), +9C chamber/+9C temperature conditioned microbiome (9C9M) and, avg(9M) which indicates the average of 9C9M and AC9M.**

| SEED L3 Classification | log2(9CAM/ACAM) | log2(9C9M/AC9M) | log2(avg(9M)/ACAM) |
|---|---|---|---|
| Nitrogen Metabolism | | | |
| Allantoin Utilization | -2.43 | NS | NS |
| Amidase clustered | -1.73 | NS | -1.14 |
| Ammonia assimilation | 2.78 | NS | NS |
| Nitrate/nitrite ammonification | 2.62 | NS | NS |
| Nitric_oxide_synthase | -1.55 | NS | NS |
| Nitrogen_fixation | -5.74 | NS | -7.99 |
| Nitrosative_stress | 1.85 | NS | NS |
| Heterocyst cyanobacteria | -2.01 | -1.64 | -8.64 |
| Sulfur Metabolism | | | |
| Sulfur_oxidation | 2.91 | NS | 2.27 |
| Taurine_Utilization | NS | NS | 1.70 |
| Glutathione as Sulphur source | 1.69 | NS | 2.51 |
| Galactosylceramid/Sulfatide_metabolism | -2.55 | NS | NS |
| Inorganic_Sulfur_Assimilation | 1.03 | NS | NS |
| Stress | | | |
| Universal_stress_protein_family | 2.41 | NS | -1.70 |
| GroEL_GroES | 1.39 | NS | NS |
| Heat_shock_dnaK_gene_cluster | 1.50 | NS | NS |
| Photo/Cyano | | | |
| Cyanobacterial_Circadian_Clock | 1.03 | NS | -4.39 |
| Myxoxanthophyll biosynthesis Cyanobacteria | 1.72 | 2.54 | -7.55 |
| Bacterial_light-harvesting_protein | NS | 4.42 | NS |
| Bacteriorhodopsin | -1.05 | NS | NS |
| PSII-type reaction_center | -1.31 | 2.85 | NS |
| Photosystem_I | 1.84 | NS | NS |
| Photosystem_II | 1.11 | NS | NS |
| Phycobilisome | 1.75 | NS | -8.89 |
| One Carbon Metabolism | | | |
| Formaldehyde_assimilation:_Ribulose_monophosphate_pathway | -1.24 | NS | NS |
| One-carbon_metabolism_by_tetrahydropterines | NS | NS | 1.98 |
| Serine-glyoxylate_cycle | NS | NS | NS |
| Soluble_methane_monooxygenase | -2.66 | NS | NS |

**SI Dataset S1 (SI_dataset1.xlsx)**

Significantly enriched MapMan4 gene ontologies for Sphagnum mapped metatranscriptome reads. Statistical significance (padj) of ontology bins was determined using Kruskal–Wallis test with multiple testing correction performed using FDR. Average log2 fold change (avg_lfc) of MapMan4 bins was determined by averaging log2 fold change across differentially expressed genes. Up/down indicates the number of genes within the bin significantly upregulated (left of slash) and downregulated (right of slash). Differential gene expression was determined using limma-voom and considered significant if the FDR corrected p-value was <=0.05. Remaining abbreviation defined in Table S7.
